# HeberFERON distinctively targets Cell Cycle in the glioblastoma-derived cell line U-87MG

**DOI:** 10.1101/2022.09.22.508971

**Authors:** Jamilet Miranda, Dania Vázquez-Blomquist, Ricardo Bringas, Jorge Fernández-de-Cossio, Daniel Palenzuela, Lidia I. Novoa, Iraldo Bello-Rivero

**Affiliations:** Bioinformatics Group for Genetic Engineering and Biotechnology (CIGB), Cuba; Pharmacogenomics Group for Genetic Engineering and Biotechnology (CIGB), Cuba; Clinical Assays Division Center for Genetic Engineering and Biotechnology (CIGB), Cuba

**Keywords:** HeberFERON, alpha interferon, gamma interferon, drug combination, U-87MG, Glioblastoma, mitotic Cell Cycle, FOXM1, PLK1, AURKB, BIRC5(Survivin), CDC20, p53, STAT1

## Abstract

**Background:** HeberFERON is a co-formulation of α2b and γ interferons, based on their synergism, that have shown its clinical superiority over individual interferons in basal cell carcinomas. In Glioblastoma (GBM), HeberFERON has shown promising preclinical and clinical results. This motivated us to design a microarray experiment aimed to identify the molecular mechanisms involved into the distinctive effect of HeberFERON compared with individual interferons.

**Methods:** Transcriptional expression profiling including a control (untreated) and three groups receiving α2b-interferon, γ-interferon and HeberFERON was performed using an Illumina HT-12 microarray platform. Unsupervised methods for gene and sample grouping, identification of differentially expressed genes, functional enrichment and network analysis computational biology methods were applied to identify distinctive patterns of HeberFERON action. Validation of most distinctive genes was performed by qPCR. Cell Cycle analysis of cell treated by HeberFERON for 24h, 48h and 72h was carried out by flow cytometry.

**Results:** The three treatments show different behavior based on the gene expression profiles. Enrichment analysis identified several Mitotic Cell Cycle related events, in particular from Prometaphase to Anaphase, that are exclusively targeted by HeberFERON. FOXM1 transcription factor network which is involved in several Cell Cycle phases and is highly expressed in GBMs is significantly down regulated by HeberFERON. Flow cytometry experiments corroborated the action of HeberFERON over Cell Cycle in a dose and time dependent manner with a clear cellular arrest since 24h post-treatment. Despite the fact that p53 was not down-regulated by HeberFERON several genes involved in its regulatory activity were functionally enriched. Network analysis also revealed a strong relation of p53 with genes targeted by HeberFERON. We propose a mechanistic model to explain HeberFERON distinctive action, based on the simultaneous activation of PKR and ATF3, p53 phosphorylation changes as well as its reduced MDM2 mediated ubiquitination and export from nucleus to cytoplasm. PLK1, AURKB, BIRC5 and CCNB1 genes, all regulated by FOXM1, also play central roles in this model. These and other interactions could explain a G2/M arrest and the effect of HeberFERON over the proliferation of U-87MG.

**Conclusions:** We proposed molecular mechanisms underlying the distinctive behavior of HeberFERON compared to individual interferon treatments, where Cell Cycle related events showed the highest relevance.

## Introduction

The molecular signaling networks underlying complex diseases limit the efficacy of single-drug treatments. Drug combination targeting multiple elements of those networks has shown advantages over one-target therapies (Zsakai et al. 2019). One such example is the synergistic effect observed by the combination of type I and II interferons (IFNs) or its combination with other cytokines (Morikawa et al. 1987, Sainz et al. 2004, Sanda et al. 2006, Thomas and Samuel 1992). On the other hand, the accelerated introduction of genome-wide technologies and network biology approaches has contributed to a faster and deeper study of drug combination mechanisms.

The co-formulation of IFNs alpha2b and gamma, HeberFERON, is produced at the Center for Genetic Engineering and Biotechnology (CIGB). This product, based on the synergism of both types of IFNs, was registered for basal cell carcinomas where the clinical superiority of HeberFERON (IFN α/γ) over individual IFN treatments was demonstrated (Bello-Rivero et al. 2013). Additionally, HeberFERON was used off-label for other types of cancer with promising results (Bello-Rivero et al. 2018). Glioblastoma (GBM) is one of the cancer entities where HeberFERON has shown clinical encouraging results (Garcia-Vega et al. 2015).

IFNs as cytokines show pleiotropic actions including antiviral and growth-inhibitory effects through several intracellular pathways from the type I and II receptors (Platanias 2005). Signaling crosstalk between IFN-α/β and -γ induce stronger responses(Taniguchi and Takaoka 2001). The understanding of the mechanisms involved in the superior results with HeberFERON, at different molecular levels, motivated the performance of a microarray experiment in the human glioblastoma derived cell line U-87MG (Clark et al. 2010).

The present work aims to shed light on the mechanism explaining the distinctive behaviors of the combination HeberFERON over the individual IFN treatments through a transcriptomic profile study. The integrative analysis of microarray data allowed us to propose a model to explain the biological outcomes and we validated cell cycle as one of the most important processes involved. This report constitutes the first high throughput experiment to investigate the distinctive HeberFERON effect in the context of GBM.

## Materials and Methods

### Reagents

Recombinant IFNs, rIFN-α2b and rIFN-γ, were produced at CIGB, Havana, Cuba. A pharmaceutical stable formulation that combines both IFN-α2b and IFN-γ (HeberFERON), was also manufactured at CIGB (Bello-Rivero et al. 2018, Bello-Rivero et al. 2013).

### Cell cultures

Human glioma cell line U-87MG (ECACC Product number 89081402, Salisbury*, Wiltshire*, UK) was maintained in complete medium MEM (Sigma, USA), supplemented with 10% Fetal Bovine Serum, 2mM glutamine and 50 µg/ mL de gentamicin (All Gibco) in a humidified atmosphere of 5% CO_2_ at 37°C. Cells were grown at a cellular density of 35 000 cells/ cm^2^. After 24 hours, cells were treated with HeberFERON at the IC50 or with equivalent quantities of rIFN-α2b and rIFN-γ using the same culture medium, and incubated for 72h. Untreated cells were included in the experimental setup.

### Cell viability assay and counting

MTT (3-(4,5-dimethylthiazol-2-yl)-2,5-diphenyltetrazolium bromide) assay was used for U-87MG viability studies. Briefly, cells were placed in 96-well culture plates (10^4^ cells/ well). After 24 h, cells were treated with 1:2 serial dilutions from 78 to 5000 total IU/mL of HeberFERON in triplicates for 72 h. At the end of treatment, 20 µL of MTT (5 mg/mL) was added to each well. The cells were incubated for another 4 h in humidified atmosphere of 5% CO_2_ at 37°C, and 100 *μ*L of 50% isobutyl alcohol-10% SDS solution were added to each well. Absorbance at 540 nm was measured and the growth inhibition ratio was calculated. Three independent experiments were performed. The half-inhibitory concentration values (IC50) were obtained from the MTT viability curves using Calcusyn software (version 2.1, Biosoft 1996-2007).

Cells were counted in a hemocytometer diluted with 0.4% of Trypan blue solution. Duplicates 175cm^2^ flasks seed with U-87MG cells (35 000 cells per cm^2^) were treated for each condition and cell counting was carried out after 72h. Number of cells is given as average ± StDev (standard deviation) in absolute number of cells or in relation to untreated control that it was considered 100%.

### Experimental design

The experiment design comprised four groups or conditions with six biological replicates each, including cells: treated with IFNα2b, treated with IFNγ, treated with HeberFERON and a control untreated group.

### RNA Purification

After 72h of incubation with IFNs medium was discarded and cells washed once with phosphate saline buffer. Cells were pickup in buffer RLT with 143mM β-mercaptoethanol and total RNA purification proceeded following the instructions of RNeasy Plus minikit (Qiagen, USA). Quality control of total RNA was carried out by spectrophotometric readings of optic density (OD) at 260 and 280nm in Nanodrop 1000 (ThermoFisher, USA) to determine concentration (> 80ng/µL) and OD260/280 ratio (1.8-2.2). Additionally, RIN (7-10) was calculated by capillary electrophoresis in a Bioanalizer (Agilent, Waldbronn, Germany). 2.5 µg (100 ng/µL) of each total RNA sample was sent to McGill University and Génome Québec Innovation Centre (Montréal, Québec, Canada) for microarray experiment in Illumina HumanHT-12 v4 Expression BeadChip platform.

### Basic Microarray Data Analysis provided by the Genome Québec Innovation Centre

The microarray experiment and a basic bioinformatics analysis were performed as a custom service at McGill University and Génome Québec Innovation Centre (Montréal, Canada). This service included quality control, preprocessing, exploratory and differential expression analysis. As a result of the quality control, none of the arrays was removed. Preprocessing included imputation for missing value using kNN algorithm, background correction and normalization using the neqc methodology described by (Shi, Oshlack and Smyth 2010). For differential expression analysis the Bioconductor Limma package (Ritchie et al. 2015) was used. Statistical tests contrasting different treatments were performed (Moderated t-tests) (Phipson et al. 2016, Zar 1999). A linear model was fit to each gene using treatment as variables. The Benjamini-Hochberg was used for FDR estimation (Benjamini et al. 2001).

### Additional Bioinformatics Analysis

To select gene expression changes that distinguish HeberFERON treatment, we compared transcription levels of treated groups against control samples. Genes with a fold change greater than 2 (|log_2_FC| >=1) and an adjusted p values of less than 0.05 (Adj p< 0.05) were considered Differentially Expressed Genes (DEGs) and used for later bioinformatics analysis. We used Venn diagrams and a scatter volcano plot that provides a summary of test statistics for DEGs.

For gene list functional enrichment analysis, the bioinformatics tools ToppGene suite (ToppFun) (Chen et al. 2009), GeneCodis (Carmona-Saez et al. 2007, Chen et al. 2009, Nogales-Cadenas et al. 2009), David (Dennis et al. 2003, Huang da, Sherman and Lempicki 2009) and BioPlanet resource were used (Huang et al. 2019). The enrichment analysis was carried against the following data sources: Gene Ontology (GO) (Ashburner et al. 2000) and biological pathways in KEGG (Kanehisa et al. 2008, Ogata et al. 1999), REACTOME (Croft et al. 2014, Fabregat et al. 2016) and Pathway Interaction Database (PID) (Schaefer et al. 2009). A cutoff value of Adjusted p value <=0.05 was set for an event to be considered significant.

Additionally, we used Gene Set Enrichment Analysis (GSEA, version 4.1.0 for windows; (Subramanian et al. 2005). Gene lists ordered by fold change were provided as input. The pre-ranked gene list option was used for REACTOME pathway database sets. We downloaded MSigdb v7.2 gmt files from: http://www.gsea-msigdb.org/gsea/downloads.jsp. The permutation-based p-value is corrected for multiple testing to produce a false-discovery rate (FDR) q-value that ranges from 0 (highly significant) to 1 (not significant). The criteria used for statistical significance was a Nominal p-value threshold of 0.05 and a False Discovery Rate (FDR) of 0.25, as recommended by the GSEA software.

BisoGenet Cytoscape plugin (Martin et al. 2010), available from Cytoscape Application Manager, was used to generate PPI networks. Venn Diagrams were generated using the web application at: https://bioinformatics.psb.ugent.be/webtools/Venn/. For the data visualization and analysis to explore brain tumors expression datasets the GlioVis Web Application was used (Bowman et al. 2017). Additional statistical analyses were performed in both: Bioconductor in R language (http://www.r-project.org/) and TIGR MeV software (The Institute for Genomic Research, USA) (Saeed et al. 2006, Saeed et al. 2003).

### Real-time PCR-based gene expression validation

Complementary (c)DNAs were obtained from 500 ng of total RNAs, using the Invitrogen SuperScript™ III First-Strand Synthesis SuperMix for qRT-PCR (Invitrogen, Carlsbang, USA) kit following manufacturer instructions. The qPCR reactions were set up in 20 µL with 300 nM of oligonucleotides (List in Table S1, *in Supplementary information*), 10 times diluted cDNAs and ABsolute QPCR SYBR Green Mixes (Thermo scientific, ABgene, UK) using three replicates per sample. The runs were carried out in a rotor equipment RT-Cycler (CapitalBio, China) using standard controls and program (Vazquez-Blomquist et al. 2012). REST 2009 (Pfaffl 2001) was used to report a Change Factor in gene levels after the treatment for 72h with IFNα2b, IFNγ and HeberFERON in relation to untreated cells after the normalization with GAPDH and HMBS as reference genes. Increases and decreases of gene levels are reported as UP and DOWN, respectively, with an associated p values (Pfaffl, Horgan and Dempfle 2002).

### Cell cycle analysis

U87MG cells were seeded in 6 well plates at 30 000cell per cm^2^ and treated with HeberFERON and IFN*α*2b or IFNγ at HeberFERON IC50 (4000 IU/mL) dose and equivalents for individual IFNs 24h after. Additional assays using 0.25XIC50 (1000 IU/mL) and 2.5X (10000 IU/mL) of HeberFERON were also carried out. Floating and adherent cells were collected by trypsinization and washed twice with PBS 24h, 48h and 72h after. Methanol (final concentration of 78%) was added to the cells, drop by drop with gently agitation, and further washed twice. Cells were then stained with 20 µg/mL of Propidium Iodide (PI) and 100 µg/mL of RNase A for 30 minutes at 37° in the dark. Finally the stained cells were analyzed and studied by flow cytometry at 488 nm (Partec CyFlow® space, GmbH, Germany; equipped with FloMax 2.9 software). Graphs of detection at FL2 channel *vs* counts were generated.

## Results

### Effects of HeberFERON over U-87MG proliferation

A viability antiproliferative study of HeberFERON over U-87MG cell line by MTT assay revealed a dose-dependent effect with an IC50 value of 4000 IU/mL (Figure S1A). Moreover, HeberFERON reduced U-87MG cell counting in a 50% using the IC50 dose. Cell counting reduction was not observed with individual IFNα2b or IFNγ at equivalent doses in HeberFERON (Figure S1B).

### Unsupervised methods revealed the four sample groups

The boxplot of microarray gene expression values shows a similar picture for all samples and the one-dimensional clustering separates samples into the four experimental groups (Fig 1A). Additionally, the 2D multidimensional scaling shows the samples for each condition grouped around each of the four corners and samples distantly located ones from the others depending on conditions (Fig 1B). The bidimensional hierarchical clustering (Fig 1C) was applied to genes showing the most variable expressions. In all cases the charts show the four experimental groups perfectly separated. In addition, the clustering shows four well distinguishable sets of genes (vertical axis) with different behaviors in each of the four conditions.

**Figure 1:**
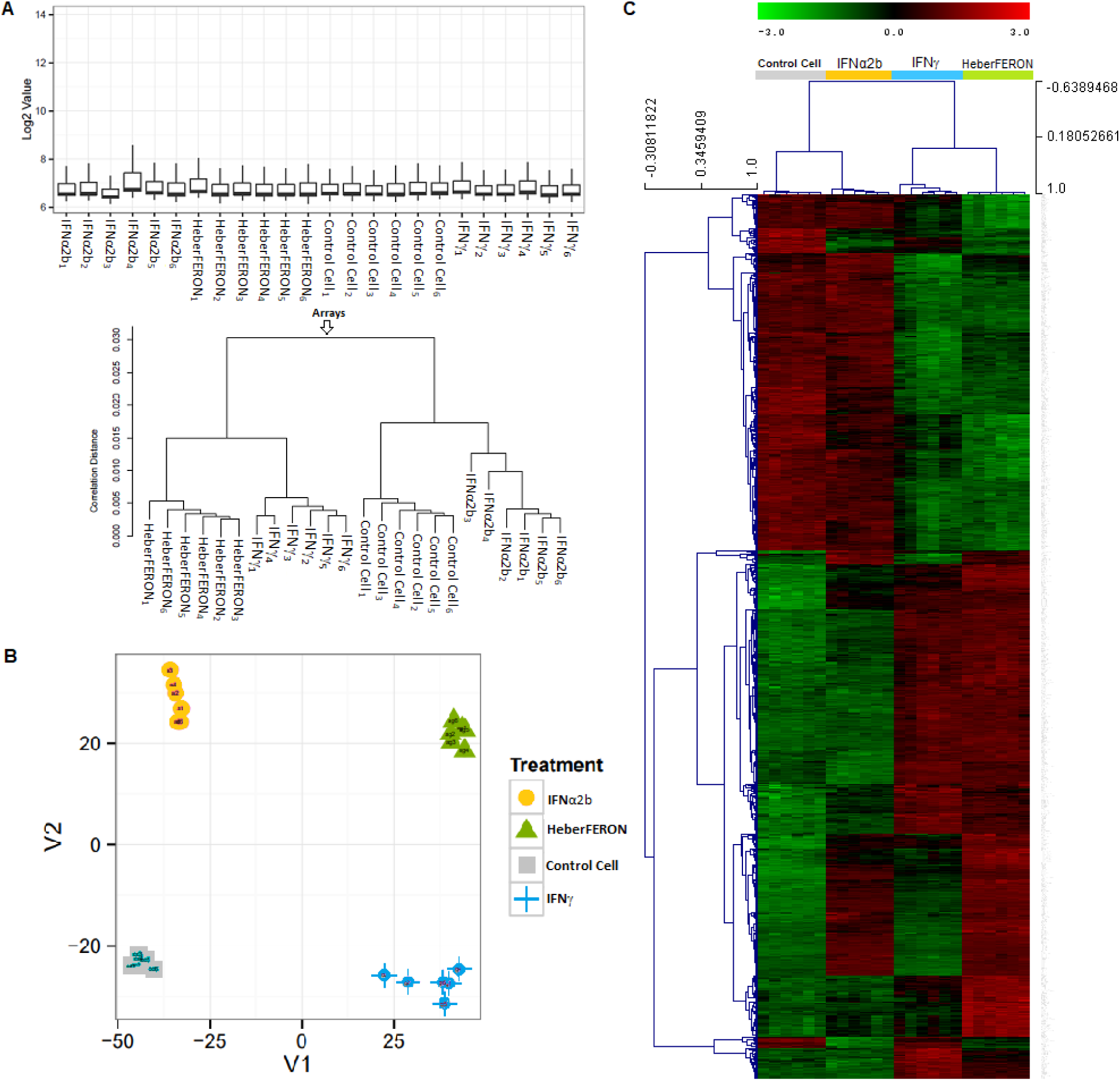
Diagnostic plots and unsupervised clustering analysis of gene expression profiles in Control cell samples and those treated with IFNα2b, IFNγ and HeberFERON. (**A**) Boxplot visualization of log2 expression values and one dimensional hierarchical clustering of all samples. (**B**) Multidimensional Scaling (MDS) of filtered data. (**C**) Two dimensional hierarchical clustering of genes with the highest fold changes. On top, samples are grouped based on similarity of their expression profiles.

### HeberFERON differentially expressed genes

Volcano plot shows the p-values *vs* FC values for each gene and treatment in the study (Fig 2A) where orange, light blue and green dots represent values for IFNa2b, IFNγ and HeberFERON treatments, respectively. DEGs are located to the left (down regulated genes) and to the right (up regulated gene). The genes with highest significant changes were predominantly from the HeberFERON treatment group. Of note it is most of DEGs by HeberFERON treatment that were down-regulated by a factor higher than 4 were no differentially expressed by individual IFN treatments. The numbers of up- and down-regulated genes for three different fold-change cut-off (2, 3, 4) are plotted in Figure 2B. IFNa2b treatment shows the smaller number of DEGs, while HeberFERON produce the highest number of DEGs. The Venn diagram in figure 2C was built with set of genes with a fold change higher than three (|FC|>3) in each of the three treatment groups. It shows 214 genes, out of a total of 563 differentially expressed by HeberFERON, that are specific for this treatment.

**Figure 2:**
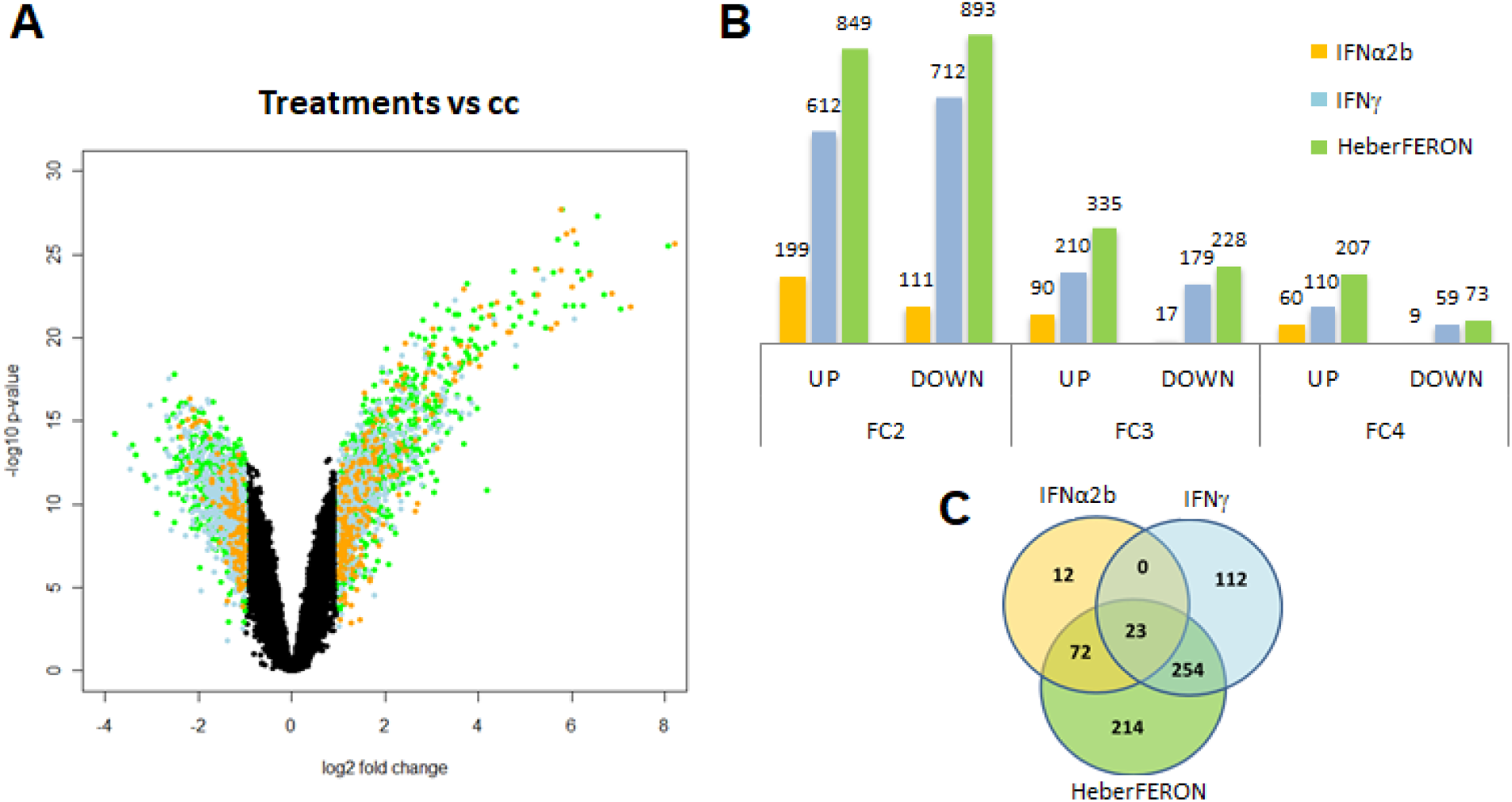
Analysis of differentially expressed genes by the action of IFN treatments. (**A**) Volcano plot visualizes significant changes between control group and treatment with HeberFERON (green points), with IFNα2b (orange points) and with IFNγ (light blue points). (**B**) Number of DEGs for different fold changes and signs of expression. (**C**) Venn diagram of sets of DEGs for the three treatment groups (|FC|>=3); IFNα2b (orange circle), IFNγ (light blue circle) and HeberFERON (green circle) groups are represented. Numbers refer to DEGs specific to one of the treatments or common to two or to the three treatments.

### HeberFERON shares some DEGs with IFNα2b and IFNγ

The enrichment analysis of Gene Ontology (GO) terms was conducted for 23 common genes to the three treatments, using David. In Table S2 we summarize the most significant biological processes (Adj p value < 0.05). The most significant processes shared by the three treatments were the defense response to virus and the innate immune response. Later on, other like antigen processing and presentation of endogenous peptide antigen via MHC class I, positive regulation of T cell mediated cytotoxicity and type I IFN signaling pathway were also included. All these are known to be activated by the action of IFNs.

### Cell cycle regulation by HeberFERON

Next we performed an enrichment analysis, using ToppGene, of sets of genes with |FC|>3 for each of the treatment. The results for each treatment were mixed and the terms reordered according to the highest level of significance (p-value). The stacked bar chart in Figure 3 shows the top 30 signaling pathways according to its significance (lower p-value). The –log_10_(p-value) for each treatment is plotted in a stacked horizontal bar. As expected the most significant pathways are those related to cytokine and IFN signaling.

**Figure 3:**
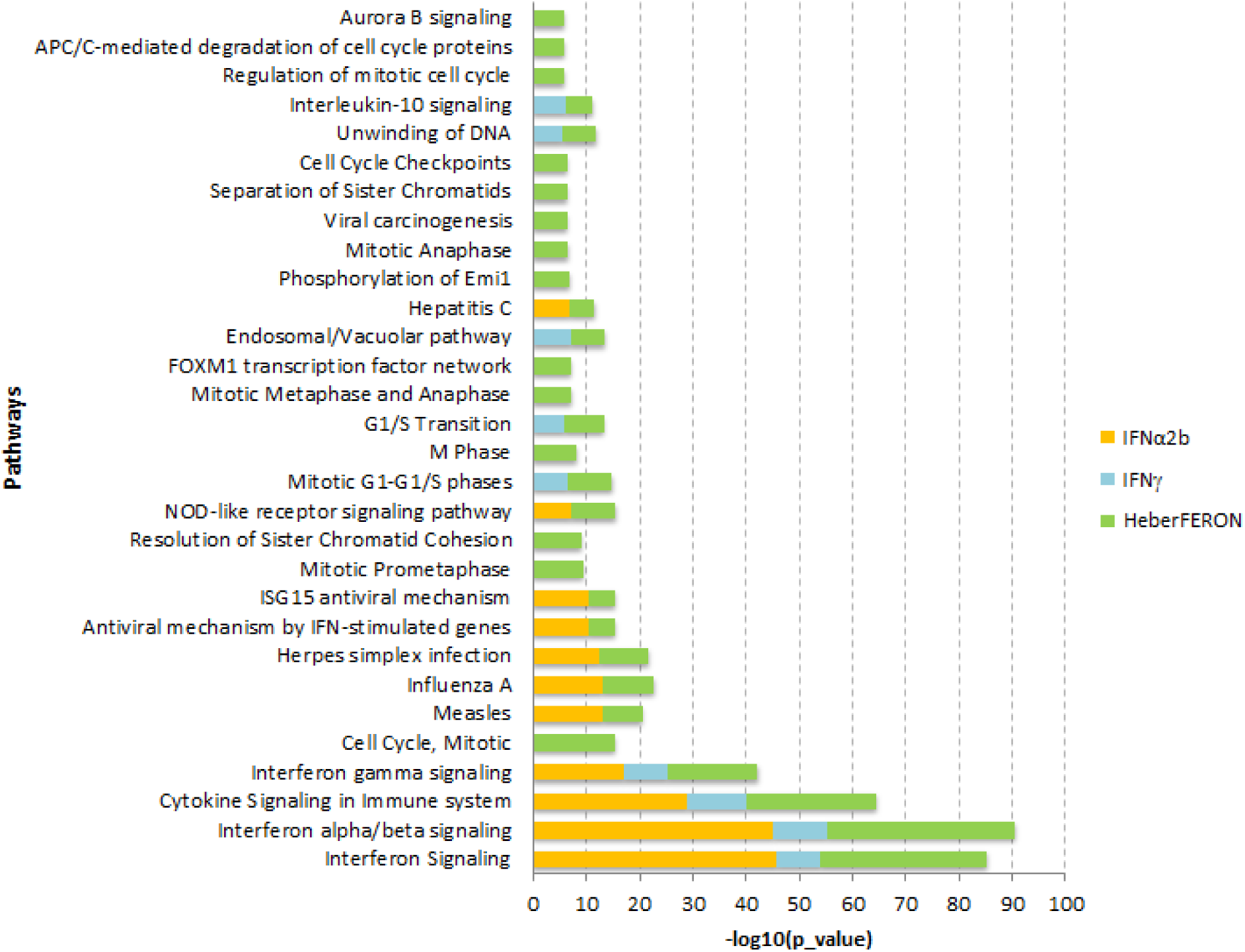
Results of ToppGene enrichment analysis for treatments. DEGs with Fold-Change three or higher were subjected to enrichment analysis. Top 30 Signaling Pathways were sorted by significance from bottom to top. Orange, blue and green color bars correspond to IFNα2b, IFNγ and HeberFERON, respectively. X axis represents the level of significance (-log10(p-value)) of enrichment scores.

The first four pathways are enriched by the three treatments. Most of the 23 DEGs common to the three treatments (Fig 2C) are involved in these four pathways. The next more significant pathway was the Mitotic Cell Cycle which together with other pathways related to cell cycle such as Mitotic Prometaphase, Resolution of Sister Chromatid Cohesion, Mitotic Metaphase and Anaphase, FOXM1 transcription factor network, Cell Cycle Checkpoints, Aurora B signaling, Phosphorylation of Emi1 and APC/C-mediated degradation of cell cycle proteins were exclusively observed in the case of HeberFERON treatment. In the case of “G1/S Transition” and “Mitotic G1-G1/S phases” pathways, IFNγ and HeberFERON treatments show similar behavior, suggesting that the IFNγ alone may also induce some inhibition of these cell cycle events.

Additionally, a comparative pathway enrichment analysis (CPEA) between HeberFERON and individual IFN treatments was conceived as follows: 1) set of DEGs by HeberFERON treatment with |FC|>3 were selected, 2) for individual IFN treatments, a less restrictive cutoff (|FC|>=2) was used, 3) the three sets of DEGs were subjected to enrichment analysis, 4) from the list of pathways enriched by HeberFERON, those enriched by either one of individual IFNs were excluded; then the resulting list of pathways could be considered as distinctively activated by HeberFERON.

Table 1 shows a list of these pathways ordered by ascending adjusted p-value. Of the 15 pathways listed, the most significant event was the “FOXM1-transcription factor network”. A network representation of this pathway is shown in Figure 4A, were nodes representing genes responsible for the enrichment are labeled and colored in orange including AURKB, BIRC5, CCNB1, CCNB2, CCNA2, CENPA, CENPF, NEK2 and PLK1. FOXM1 is highly expressed in GBM (Figure 4B) and genes regulated by this transcription factor have also the highest expression in GBM compared to others brain tumors (Figure S2). Together with mitotic cell cycle related pathways already found by ToppGene enrichment analysis, we found “p53 activity regulation” as a new enriched event. In spite of p53 was not differentially expressed by HeberFERON, several genes participating in its signaling including cyclins (CCNA2, CCNE2, CCNB1, CCNB2, CCND2) were down-regulated by HeberFERON.

**Table 1:** List of the top BioPlanet significant pathways regulated by HeberFERON as the result of CPEA.

| Pathway_id | Description | Adj p value | Genes |
| --- | --- | --- | --- |
| bioplanet_1345 | FOX M1 transcription factor network | 1.87E-05 | AURKB, BIRC5, CCNA2, CENPF, FOXM1, CENPA, CCNB2, CCNB1, PLK1, NEK2 |
| bioplanet_592 | Aurora B signaling | 1.03E-04 | STMN1, AURKB, CDCA8, BIRC5, CENPA, NCAPG, INCENP, KIF23, KIF20A |
| bioplanet_1622 | Phosphorylation of Emi1 | 3.82E-04 | FBXO5, CDC20, CCNB1, PLK1 |
| bioplanet_67 | Stathmin and breast cancer resistance to antimicrotubule agents | 1.40E-03 | HIST1H4C, STMN1, CREB1, OAS1, USP18, HLA-B, STAT1, GBP1, CXCL10, PRKACB, IRF1, IFIT2, KAT2B, IFI6, TAP1, ISG15, EIF2AK2, RNF14, CCNB1, STAT2, PSMB9, IRF9 |
| bioplanet_1327 | Polo-like kinase 1 (PLK1) pathway | 1.79E-03 | FBXO5, CDC20, PRC1, INCENP, CCNB1, SPC24, PLK1, KIF20A |
| bioplanet_1757 | Hypertrophy pathway | 5.99E-03 | IFRD1, ATF3, VEGFA, CYR61, IL1A |
| bioplanet_1391 | APC/C activator regulation between G1/S and early anaphase | 7.36E-03 | FBXO5, CDC20, UBE2C, CCNB1, PLK1 |
| bioplanet_1363 | Delta Np63 pathway | 1.09E-02 | NRG1, FASN, RRAD, TOP2A, CCNB2, IL1A, RAB38 |
| bioplanet_235 | Cytokine-cytokine receptor interaction | 1.31E-02 | TNFSF13B, CSF3, IL11RA, INHBE, CXCL5, TNFSF10, CSF1R, CXCL10, CCL2, IL20RB, CCL20, CXCL2, LEP, VEGFA, IL1A, IL6, IL24, TNFRSF10D, CCL8 |
| bioplanet_884 | APC/C- and Cdc20-mediated degradation of Nek2A | 1.47E-02 | MAD2L1, CDC20, UBE2C, CCNB1, NEK2 |
| bioplanet_1511 | Cyclin A/B1-associated events during G2/M transition | 1.49E-02 | CCNA2, CCNB2, CCNB1, PLK1 |
| bioplanet_195 | p53 activity regulation | 1.92E-02 | GADD45A, KAT2B, SESN2, CCNA2, CCNE2, SHISA5, CCNB2, PMAIP1, GTSE1, CCNB1, CCND2 |
| bioplanet_1575 | Kinesins | 1.95E-02 | KIF22, KIFC1, KIF11, KIF23, KIF20A |
| bioplanet_1139 | MicroRNA regulation of DNA damage response | 2.19E-02 | CREB1, MCM7, GADD45A, CCNE2, CCNB2, PMAIP1, CCNB1, CCND2 |
| bioplanet_1409 | Mitotic prometaphase | 2.82E-02 | ZWINT, CDC20, CENPF, CENPA, INCENP, PLK1 |

**Figure 4:**
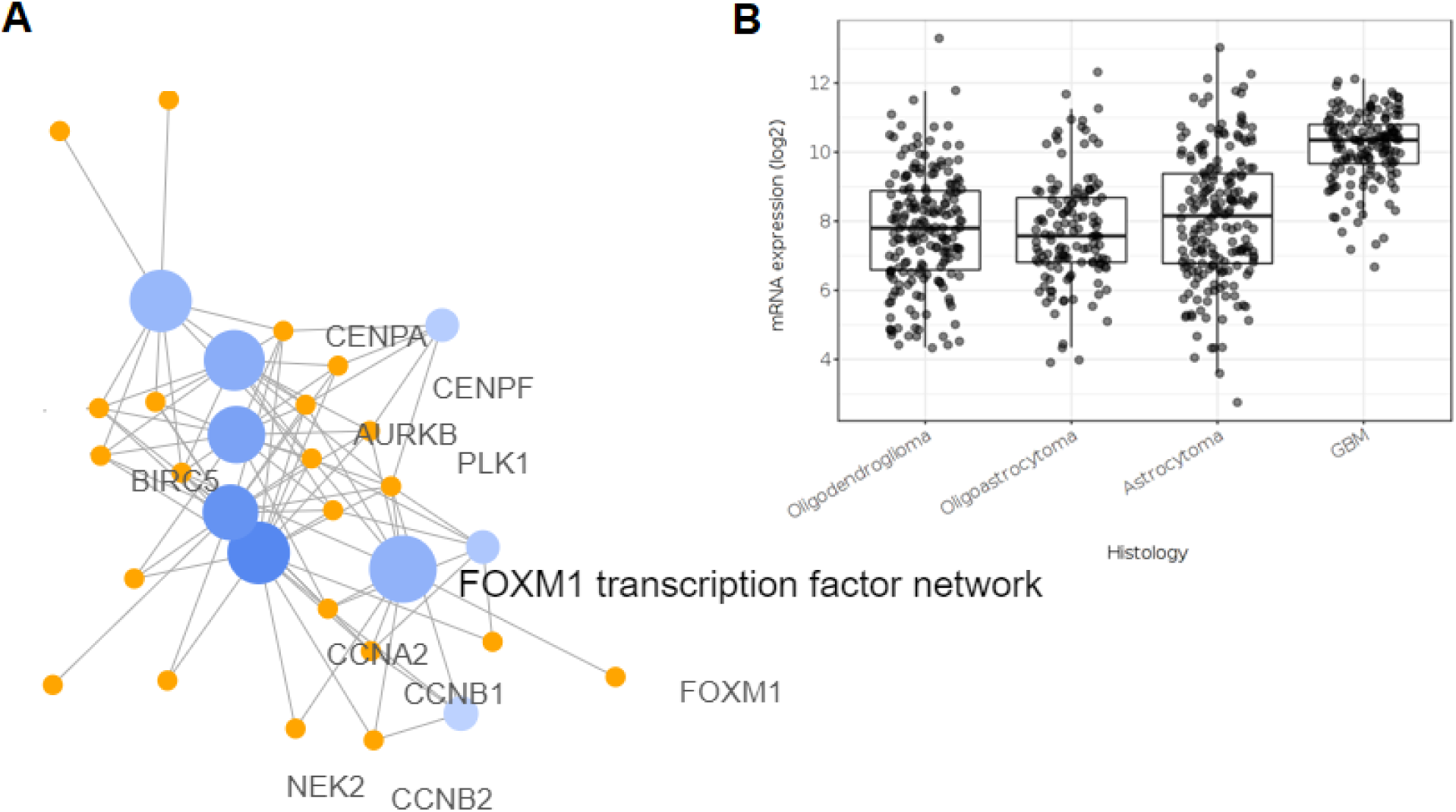
FOXM1 transcription regulation and its expression in brain tumors. (**A**) Network of BioPlanet enriched pathways (blue nodes) and genes (orange nodes) using GeneCodis. Nodes belonging to the FOXM1 transcription factor network are labeled. (**B**) FOXM1 expression pattern in brain tumors (oligodendrogliomas, oligoastrocytoma, astrocytoma and GMB) datasets using GlioVis application.

From the 15 enriched pathways we identified 75 DEGs. The heatmap in Figure 5A groups the expression profiles in four main clusters of genes (vertically) and the four samples (horizontally). The first two gene clusters (orange and blue lines) contain genes up regulated by IFNα2b and IFNγ, respectively. HeberFERON up-regulates most genes in both clusters. The third cluster (magenta line) contains a few chemokines while the fourth cluster (green line) contains genes involved in several enriched Cell Cycle events which are strongly down-regulated by HeberFERON. Figures 5B, 5C and 5D show the expression profiles of genes related to the first three events listed in Table 1 (“FOXM1 transcription factor network”, “Aurora B signaling” and “Phosphorylation of Emi1”). These genes belong to the forth cluster in Fig 5A.

**Figure 5:**
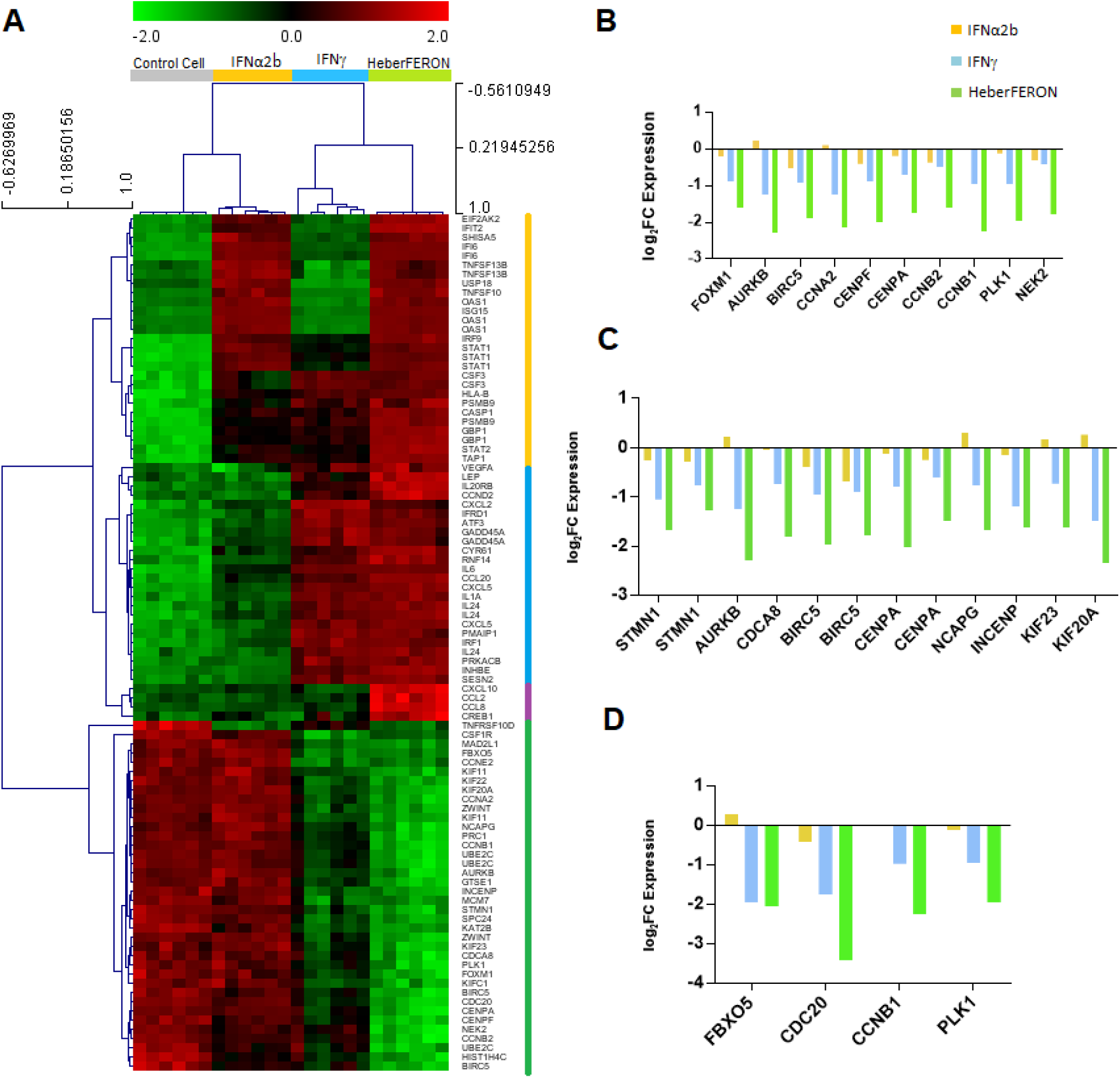
Gene expression analysis of genes resulting from CPEA of BioPlanet pathways. (**A**) Heatmap of all genes resulting from CPEA analysis in Control cell samples and those treated with IFNα2b, IFNγ and HeberFERON. The four colors in the vertical line to the right of the heatmap identifies four different gene clusters: first cluster (orange) groups genes up-regulated by IFNα2b and HeberFERON; second cluster (light blue) groups genes up-regulated by IFNγ and HeberFERON; third cluster (magenta) groups a small set of genes upregulated by HeberFERON and four cluster (green) groups genes down-regulated by the HeberFERON treatment. (**B**), (**C**) and (**D**) log2FC expression value of genes in the top three enriched pathways resulting from CPEA: “FOXM1 transcription factor network”, “Aurora B signaling” and “Phosphorylation of Emi1”, respectively.

As an additional analysis we applied Gene Set Enrichment Analysis (GSEA), a method that does not require any filtering by a threshold. As input we provided lists of all genes in the microarrays ranked by fold change. The method identifies enriched terms in the top or bottom of the ranked list. In supplementary Figure S3, GSEA results for the effect of HeberFERON and IFNγ are provided. Pathways are ranked by the Normalized Enrichment Score (NES) of HeberFERON treatment. The top 12 pathways in the list are related to Cell Cycle and clearly more down-regulated by the HeberFERON treatment (Figure S3A). Figures S3B and C show the enrichments plots and most relevant genes from Core enrichment set of events only enriched by HeberFERON: “Deposition of new CENPA containing nucleosomes at the centromere” and “Kinesins”. Additionally, in Figure S4 the expression levels of genes involved in Cell Cycle are shown; most of them are more down-regulated by HeberFERON than by individual IFN treatments. These results reinforce Cell Cycle related events as distinctively targeted by HeberFERON in comparison to individual IFN treatments.

### Network analysis of genes targeted by HeberFERON

Figure 6 shows a network of DEGs by HeberFERON plus p53 composed by 230 connected nodes. p53 was added to the network because p53 activity regulation was one of the enriched events by CPEA. Of note the node with the highest degree is that representing tumor suppressor p53, suggesting a role of this protein to mediate HeberFERON action.

**Figure 6:**
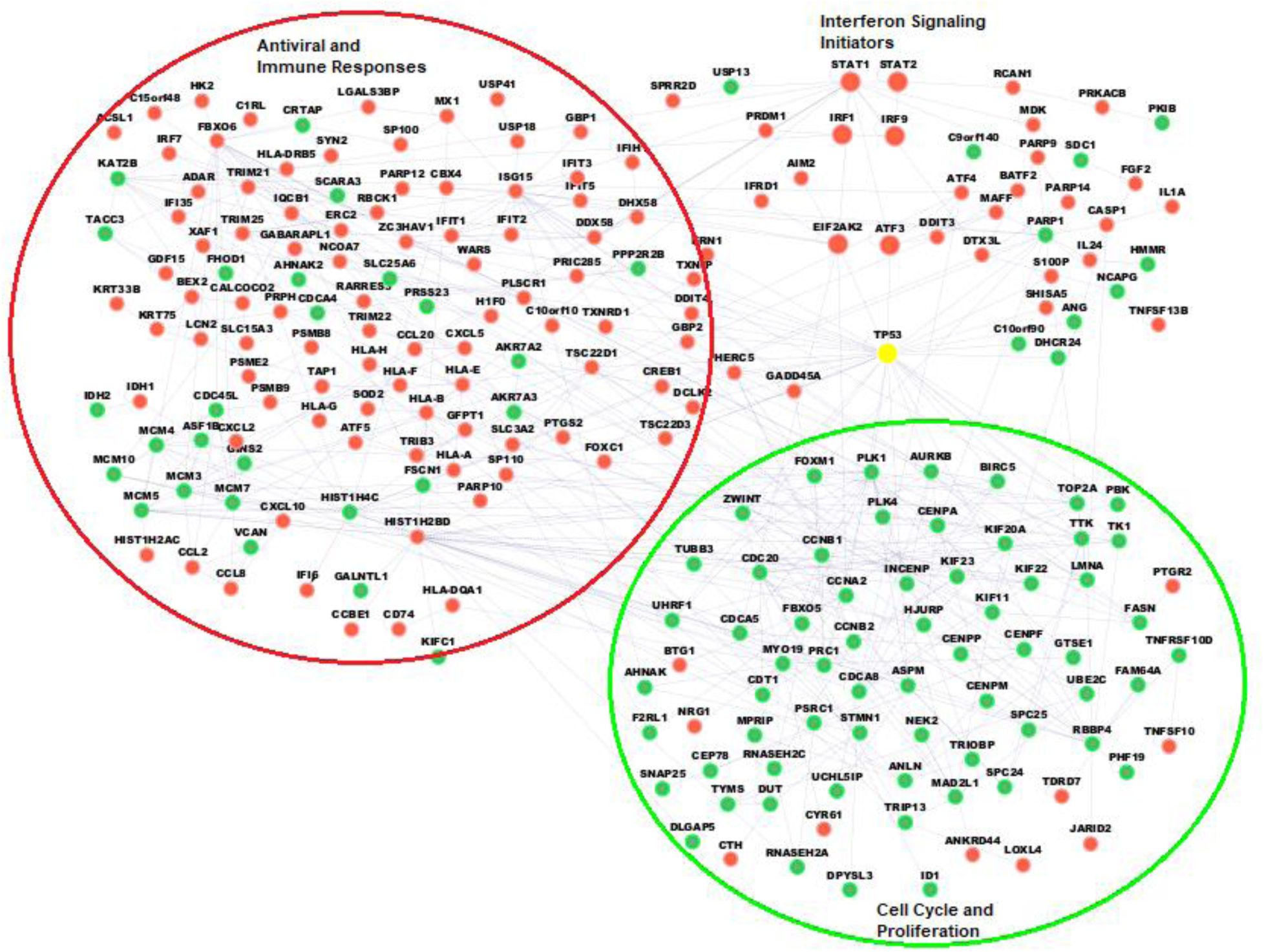
Network of DEGs by HeberFERON treatment plus p53. A total of 230 nodes are represented, the rest of the 563 DEGs were not connected or were part of small subnetworks. Red and green colors represent up-regulated and down-regulated DEGs, respectively. In yellow the node representing the p53 tumor suppressor, the one with the higher degree. The network was generated by BisoGenet CytoScape application.

This network shows the high interconnection between DEGs from the STATs, but also subsets of genes participating in common pathways for IFN treatments as IFN signaling, antiviral and immune responses. P53 encoding gene (TP53) looks as that high degree node from where the signaling transduction cascade connect to biomarkers participating in cell cycle events (PLK1, AURKB, ZWINT, CCNA2), proliferation (TOP2A, TK1, PBK, TTK) or replication (MCM family members). Most of these biomarkers are down-regulated by HeberFERON.

These effects are closest shown when we built a network with only the 75 genes resulting from the CPEA plus p53 (Figure 7). A 50 connected nodes network is composed of two main sub-networks. The first sub-network is composed of key mediators of IFN Signaling, including STAT1, STAT2 and IRFs genes, predominantly up-regulated by HeberFERON. A second subnetwork included genes that are involved in Cell Cycle Mitotic events predominantly down-regulated by HeberFERON. Here, it is evident P53 encoding gene (TP53) is located in the interface between both sub-networks with the highest degree nodes connection to both subnetworks.

**Figure 7:**
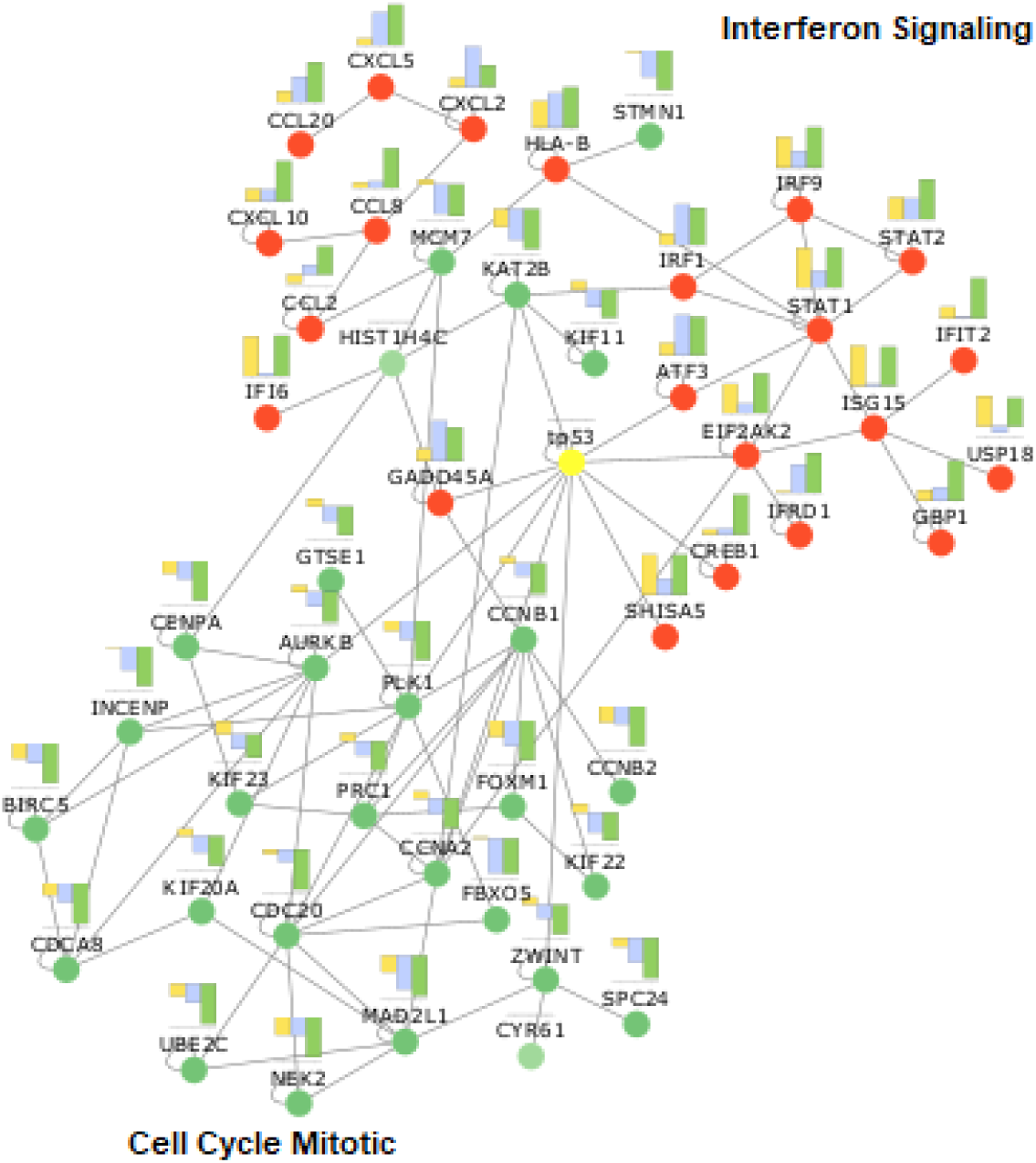
Network of the interconnected genes (50 of 75) selected from the CPEA plus p53. Red and green colors represent up-regulated and down-regulated DEGs, respectively. In yellow the node representing p53 tumor suppressor. For each node a bar chart shows the expression levels of each gene in the three experimental groups: orange, blue and green color bars correspond to IFNα2b, IFNγ and HeberFERON, respectively. The network was generated by BisoGenet CytoScape application.

A PubMed enrichment analysis using ToppFun identified a signature of poor prognosis in Proneural subtype of GBM (PMID: 22242177). The genes involved in this signature are closely related to STAT1 which is up-regulated by HeberFERON. The proximity of IFIT1, IFIT3, ISG15, MX1, STAT1 and USP18 signature genes in the interaction network is evidenced in Figure 6. In supplementary Figure S5 we show the expression levels of genes belonging to this poor prognosis signature for each IFN treatment.

### Validation of regulated cell cycle gene expression by qPCR

We performed qPCR validation of a set of genes participating in Cell Cycle regulation, among them FOXM1 and members of its regulatory network and prometaphase proteins, PLK1, AURKB, BIRC5, CCNB1, CENPA, CENPF and ZWINT. We also included some other genes encoding proteins participating in the spindle checkpoint as CDC20, BUB1, BUB1R and CENPE. All of them showed a higher expression decrease with HeberFERON treatment (Figure 8).

**Figure 8:**
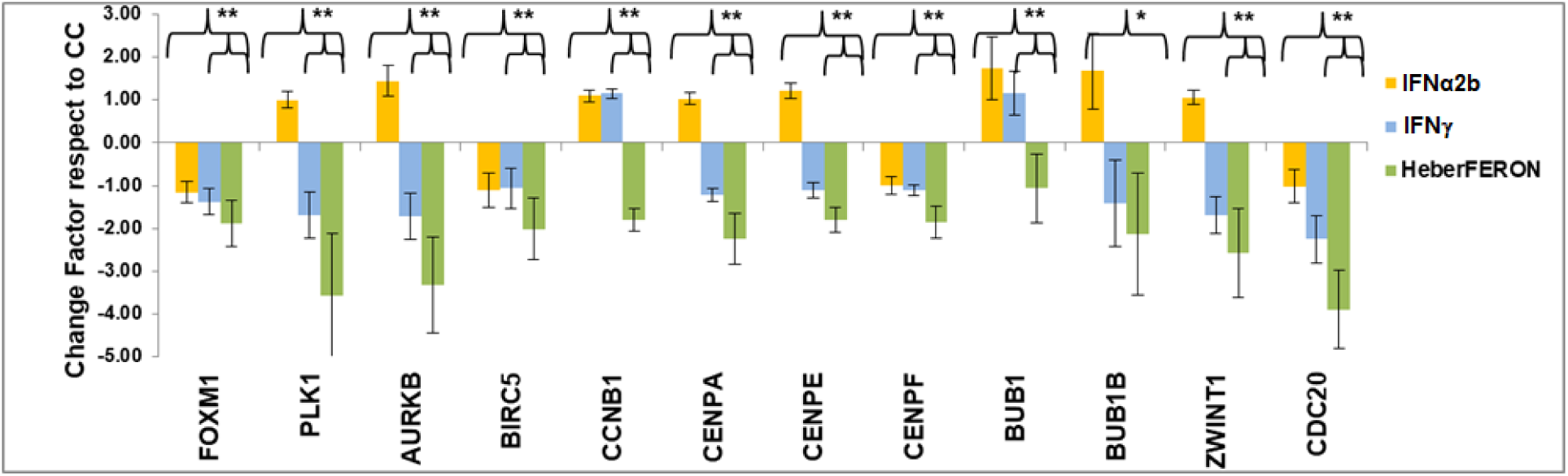
Fold change of transcript levels for genes involved into Cell Cycle regulation by qPCR. Fold change in transcript levels with respect to untreated controls are shown for the treatment with IFNα2b, IFNγ or HeberFERON and the standard error associated with the measurements. The statistically significant differences according to REST2009 (p <0.05) are shown with asterisks (*) for the comparisons indicated with parentheses.

### Cell Cycle analysis

We tested if HeberFERON modulates cell cycle dynamics in U-87MG cells, in comparison to individual IFNα2b and IFNγ (Figure 9A). HeberFERON induced an S-G2/M cell cycle arrest after 72h of treatment using the IC50 dose (27.3 % of cells in these phases compared to 8.9 % for untreated culture), whereas IFNα2b and IFN induced some arrest but in a lower percentage of cells (17.5 % and 14.8%, respectively). This arrest is observed as early as 24h after treatment (Figure 9B), a time where the cycle dynamic is highly hit. This effect is dose and time dependent as it is shown in Figure 9C. The extension of the effect at a dose of 0.25XIC50 (1000 IU/mL) is barely observed but S-G2/M arrest can be seen at IC50 (4000 IU/mL) and 2.5XIC50 (10000 IU/mL), with a higher impact at 72h with 10000 IU/mL dose of HeberFERON.

**Figure 9:**
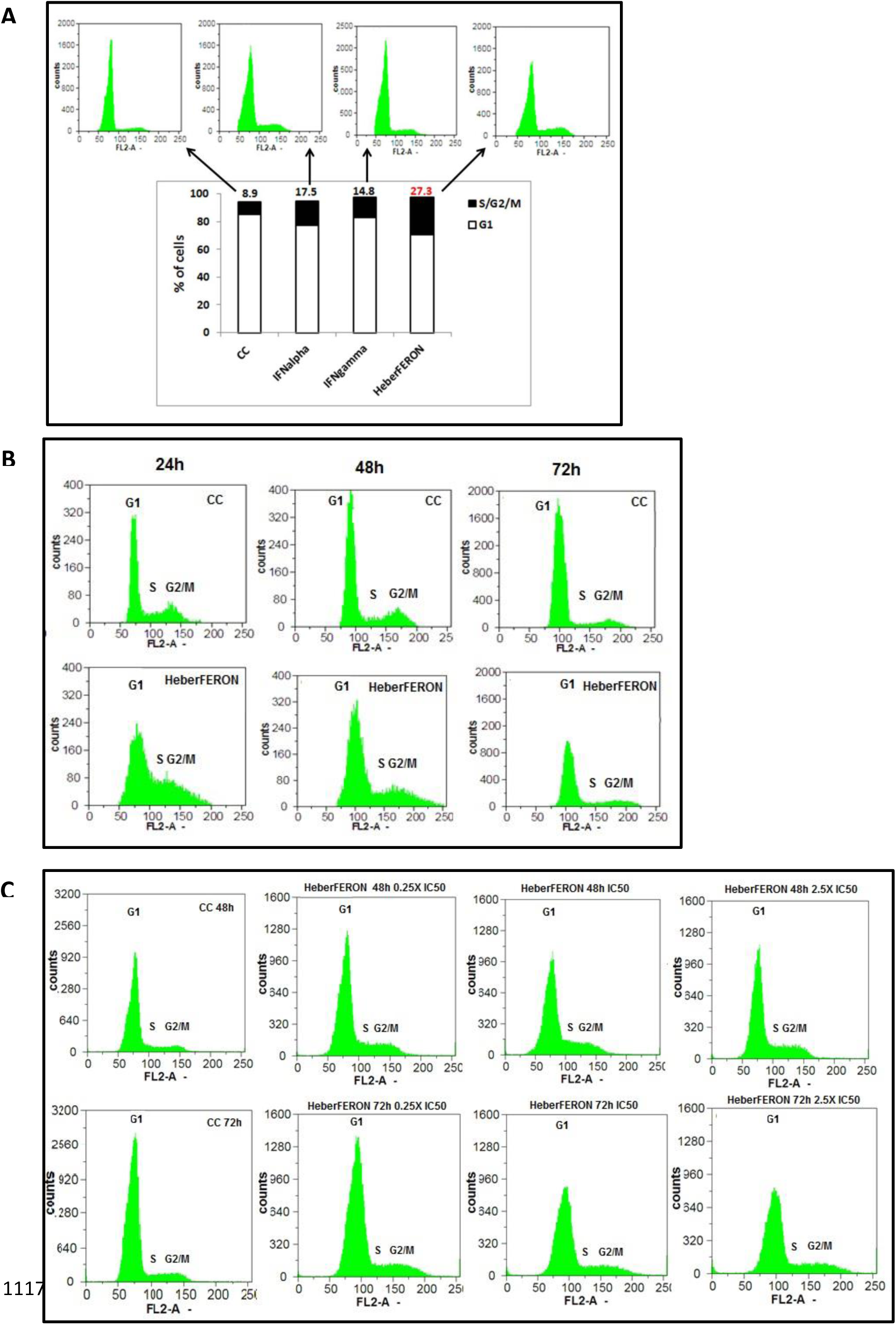
S-G2/M cell cycle arrest in U87MG cells by HeberFERON, dose and time dependence. (**A**) U87MG cells were pre-incubated with IC50 of HeberFERON and equivalent individual IFNα2b and γ for 72h and cell cycle analysis by flow cytometry. The figure shows the counts vs PI staining in FL2 channel in the untreated control (CC) in comparison to treatment with IFNα2b, IFNγ or HeberFERON in relation to the % of cells in phase G1 or S/G2/M; Cell cycle analysis was carried out (**B**) at 24h, 48h and 72h of cell treatment with HeberFERON (IC50=4000 IU/mL) and (**C**) at 48h and 72h treating cells using 1000 (0.25XIC50), 4000 (IC50) and 10000 (2.5XIC50) IU/mL of HeberFERON.

## Discussion

Glioblastoma is one of the most aggressive types of malignant central nervous system tumors, with a rapid infiltrative growth rate and provoking a heterogeneous disease (Ostrom et al. 2021). Despite various therapeutics have been assayed, there have been limited advances in the overall survival (OS) rate of GBM (Janjua et al. 2021). Due to this void in real effective therapy new approaches allowing the increase of OS and/or improving patients’ quality of life are welcome.

The use of IFNs in cancer therapy, including glioma has been reported before with non-homogeneous results (Dapash et al. 2021, Janjua et al. 2021), however the HeberFERON, has shown promising results in clinical trials (Bello-Rivero et al. 2018). The understanding of its molecular mechanism of action could guide us to redesign therapies, suggests new formulations and combination with other anti-cancer drugs. Here, we report the first high throughput transcriptomic analysis of the model human glioblastoma cell line U-87MG treated with HeberFERON, IFNα2b and IFNγ for 72h. As it was previously found by Tan *et al*, 2005 and Sanda *et al*, 2006, our experiment showed the combination of type I and type II IFNs modulates a much larger number of genes than either individual IFNs (Sanda et al. 2006, Tan et al. 2005).

We showed how multidimensional scaling was capable to distinguish samples in each group, but there are common processes and/or genes between groups related to very well established IFN functions in the antiviral, immune and inflammatory responses. Previous studies evidenced type I and type II IFNs combination had a functional consequence up-regulating ISGF3 components, enhancing expression of ISRE and GAS-containing genes associated with a direct antiviral state (Tan et al. 2005).

Tan *et al* also reported 26% of the probe sets with differential modulation corresponded to components involved in antigen presentation and processing, immune cell recruitment or complement system function. Several interferon-stimulated genes (ISGs) as those encoding for 2’-5’ OAS, RNaseL, PKR or IRF9 also participate in the antiproliferative effects (Bekisz *et al*, 2010). These functions have been exploited in the treatment of viral and neoplastic diseases. In a context of a brain tumor, immune response regulation could also contribute to the overall effect of this product (Dapash et al. 2021).

Enrichment analysis showed similarities and differences between the three treatments with a remarkably distinctive behavior of HeberFERON targeting a range of Cell cycle events and more specifically, mitotic cell cycle. Meta-analysis from several gene expression studies in GBM has previously concluded that Mitosis is one of the most relevant biological events in this complex disease (Horvath et al. 2006).

Cell Cycle is a highly regulated process at transcription level, by phosphorylation or protein localization changes and degradation. From G1, cells increase in size, copy DNA and duplicate centrosomes in S and prepares in G2 to divide in Mitosis. Surveillance mechanisms through Cyclins and Cyclin-dependent-kinases (CDK) ensure proper timing of cellular events at G1/S, G2/M and spindle-assembly Checkpoints. Later on, the transition toward Prophase, Prometaphase, Metaphase, Anaphase and Telophase mitosis stages occurred by ordered protein degradation by SCF (Skp1/Cullin/F-box) and APC/C protein complexes (Poon 2016).

Along these stages chromosomes primarily condense and centrosomes begin to separate in Prophase; mitotic spindle forms and the interaction of microtubules with the spindle and kinetochore protein complexes at the centrosomes, in Prometaphase, will permit the future separation of sister chromatids to opposite poles of the cells. In Metaphase/Anaphase transition chromosomes should be bi-oriented and Spindle Checkpoint will ensure the formation of this configuration. Spindle poisons leading to cancer cells mitotic arrest have encouraged the search for new inhibitors. There are promising ongoing trials for GBM targeting therapy with G2/M inhibitors, including inhibitors of Aurora kinases, PLK1, Survivin, BUB1 and BUBR1 (Castro-Gamero et al. 2018).

Among the events targeted only by HeberFERON, are the Mitotic Prometaphase, Metaphase and Anaphases as well as Cell Cycle Checkpoints. As part of these events we found genes with the highest down-regulation by HeberFERON. This is the case of those encoding for Prometaphase proteins PLK1, AURKB, BIRC5/Survivin, CCNB1, CENPA, CENPF, ZWINT1 and proteins participating in the spindle checkpoint as CDC20, BUB1, BUB1R and CENPE. Moreover, FOXM1 transcription factor network was the top enriched pathways by HeberFERON when we applied the CPEA, with a significant down-regulation of genes involved. This transcription factor plays an essential role in mitotic progression in general (Wang et al. 2005) and it is critical in GBM development and progression becoming into an attractive drug target (Wang et al. 2015). If we looked at the expression of FOXM1 in multiple samples from four different types of brain tumors in GlioVis data portal, it is evident the highest expression in GBM. The use of HeberFERON may delay the progression of GBMs and also contribute to reduce the resistance to Temozolomide (TMZ) treatment. TMZ is the standard chemotherapy for GBM since 2005 (Singh et al. 2021) and combined with radiotherapy, it added two-month increase in OS as average (Stupp et al. 2005). GBM therapy failure can be due to TMZ resistance, enhanced by CXCL12/CXCR4 promotion of migration of GBM cells by up-regulating FOXM1. Thus FOXM1 silencing can partially reverse this resistance (Wang et al, 2020). Regulation of mRNAs expression by FOXM1 permits accumulation of cyclin B1 during G2 and decrease after mitosis. FOXM1 also controls AURKB, CENPA, CENPF, NEK2, PLK1 and Survivin. Most of these genes are commonly overexpressed in different types of human cancer, including GBM (Alafate et al. 2019, Cheng et al. 2012, Tong et al. 2019, Zeng et al. 2007) which it was also shown in GlioVis portal analysis of brain tumor samples.

AURKB and Survivin, together with Borealin and INCENP, composed the chromosomal passenger complex that localizes to the kinetochores and chromosomes during early mitosis and functions in microtubule–kinetochore interactions, sister chromatid cohesion, and spindle-assembly checkpoint (Honda, Korner and Nigg 2003).

Aurora kinases regulate different aspects of cell division. Aurora kinase B (AURKB) is present in the centromeres in Prophase and Metaphase and it is located in the central mitotic spindle, being crucial for the segregation of chromosomes and cytokinesis (Zeitlin, Shelby and Sullivan 2001). AURK A and B expressions and kinase activities are elevated in a variety of human cancers and associated with high levels of proliferation and poor prognosis (Sankhe, Prabhu and Khan 2021). AURKB was proposed as a prognosis marker for GBM from a study that showed an overexpression in 25 GBMs samples and a correlation of its expression levels with survival (Zeng et al. 2007). Small molecules inhibitors of Aurora kinases A and B interfere with the centrosome function during mitosis and disrupt the assembly point of the mitotic spindle resulting in polyploidization and apoptosis of proliferating cells (Li et al. 2009, Tang et al. 2017). The strong down-regulation of AURKB by HeberFERON may contribute to increase the OS of GBM patients.

Survivin (BIRC5) acts as a suppressor of apoptosis and plays a central role in cell division. It is expressed in the G2/M phase, is located in the mitotic spindle interacting with tubulin and also plays a role in the regulation of mitosis. It was also reported to be located in the centromeres influencing the stability of the kinetochore-microtubule junction and in the control signal of the mitotic spindle, physically interacting with AURKB (Beardmore et al. 2004). It is also highly expressed in most cancers; in some subtypes, it has a prognostic value related to antineoplastic resistance and radiotherapy as occur for Cisplatin (Zaffaroni and Daidone 2002). Several studies have shown that the inhibition of Survivin reduces tumor growth, increasing apoptosis and sensitizing the tumor to different chemotherapeutic agents (Cheung et al. 2013, Mita et al. 2008). Hence, the inhibition of BIRC5 by HeberFERON suggests its possible combination with various chemotherapeutic (vincristine, cisplatin, bortezomib, tamoxifen) to avoid resistance. This protein has been shown to be essential for proliferation but is not required for the survival of normal cells (Li, Hu and Li 2018). According to Beardmore et al, (2004) BIRC5 regulation appeared to be linked to p53 protein.

PLKs belong to a family of Serine-Threonine kinases that play key roles in the control of the cell cycle and in the response to DNA damage. In G2/M checkpoint, cells with damaged DNA are prevented to enter Mitosis. Cyclin B-CDK1 complex activity is essential at this point and a feedback amplification loop is established among Aurora kinase A, PLK1 and CDK1 by phosphorylation. Many studies have shown PLK1 inhibition lead to death of cancer cells by interfering with multiple stages of mitosis (Danovi et al. 2013). PLK1 mRNA expression strongly correlated with WHO grades, KPS and the recurrence of tumors of patients with gliomas. Down-regulation of PLK1 at the level of mRNA and protein was able to inhibit growth, induce the arrest of the Cell Cycle in G2/M and increase glioma cells apoptosis (Pezuk et al. 2013). PLK1 promotes the translocation of cyclin B into the nucleus during prophase and initiates the cycling by activating phosphatase CDC25 and inactivating WEE1/MYT1 kinases. Activated cyclin B–CDK1 stimulates the activity of APC/C CDC20 through phosphorylation of several subunits of APC/C and CDC20 but APC/C is also phosphorylated and activated by PLK1. Besides the role of Cyclin B-CDK1 complex in G2/M checkpoint, its activity also contributes to the inactivation of the mitotic spindle checkpoint. The loss or low expression of Cyclin B1 (CCNB1) causes a deficient binding between the kinetochores and the microtubules, defects in the alignment of the chromosomes and delays the entry to Anaphase (Chen et al. 2008). HeberFERON causes a significant decrease in CCNB1 that distinguishes it from individual IFNs effects. This could be contributing to Anaphase entry impairment and Cell Cycle arrest.

Moreover, GSEA results show CENPA function at the centromere and Kinesins could also have roles in HeberFERON molecular mechanism. CENPA is required for kinetochore recruitment of all other kinetochore components. Phosphorylation of CENPA by AURKB plays an important role in cytokinesis (Zeitlin et al. 2001), Ser7 phosphorylation by AURKA is required for concentration of AURKB at centromeres and for the function of kinetochore (Kunitoku et al. 2003). HJRUP interact with CENPA and it is required for its centromeric assembly and deposition (Foltz et al. 2009, Zasadzinska et al. 2013). Here, CENPA, HJURP and the inner kinetochore components CENPM and CENPP decreased expressions by HeberFERON. Kinesins are crucial in different stages of cell division. Particularly, KIF23 and KIF20A have a major role in citokinesis (Rath and Kozielski 2012). KIF23 is responsible for bundling and stabilizing microtubules, it requires INCENP for its recruitment (Zhu, Bossy-Wetzel and Jiang 2005) and it is regulated by CDK1, Aurora B and PLK1 (Lee et al. 1995, Liu et al. 2004). Its down-regulation was shown to suppress glioma proliferation, being proposed as a potential therapeutic target for GBM (Takahashi et al. 2012). PLK1 directly phosphorylates KIF20A and regulates its motor properties, and at the same time, KIF20A seems to be essential for normal localization of PLK1 to the central spindle (Neef et al. 2003). HeberFERON down-regulates a set of Kinesins including KIF23, KIF20A, KIF11, KIF22 and KIFC1 that could affect its motor functions.

Spindle checkpoint is activated by either the presence of unattached kinetochores or the absence of tension between paired kinetochores. Kinetochores couple sister chromatids to dynamic microtubules during congression and anaphase; this allows their separation and partition to the daughter cells (Musacchio and Desai 2017). Unattached kinetochores attract several components of the checkpoint sensors (including BUB1, BUBR1, CENPE and MAD2), catalyzing the formation of mitotic checkpoint complexes (in the outer kinetochore), resulting in the inhibition APC/C-CDC20.

It is also evident HeberFERON induced a marked decrease of genes encoding checkpoint mitotic complex components as BUB1, BUB1R, CENPE, CENPF and CDC20. Morrow et al (2005) suggested checkpoint is composed by one arm dependent on BUB1 and the other on AURKB, both converging on the mitotic checkpoint complex (Morrow et al. 2005). Furthermore, BUB1R kinase is regulated by CENPE and at the same time it regulates the proteolytic machinery APC/C-CDC20. Knockdown of BUB1B/BUBR1 inhibited expansion of brain tumor–initiating cells isolates, both *in vitro* and *in vivo*, without affecting proliferation of human neural stem cells or astrocytes (Ding et al. 2013). These results point this protein as the top-scoring glioblastoma lethal kinase. HeberFERON then hit multiple transcripts for proteins that are important to pass this control point in cell cycle.

Zhou et al (2019) found CDC20, TOP2A and PBK to be highly up-regulated in glioblastoma samples compared with healthy tissue. These genes were all identified as hub genes in DEGs network and inhibited by HeberFERON. A Bisogenet network was built with the immediate neighbors of CDC20, TOP2A and PBK containing 275 highly connected nodes; CDC20, P53 and TOP2A showed the highest degree, in that order. TTK was identified as the most up-regulated gene encoding protein kinases in glioma stem-like cells, it was essential for clonogenicity and tumor propagation correlating with poor prognosis in GBM patients (Wang et al. 2018). HeberFERON also down-regulates TTK. These genes together with ZWINT, HJURP, CENPA and several kinesins are connected in a network from the cascade initiators STAT1&2, passing through PKR and ATF3 nodes, all these transcription factors highly increased with HeberFERON.

In correspondence with the above interpretation, cell cycle FACS analysis showed a dose and time dependent effect of HeberFERON over the process, with a clear effect of arrest since 24h of treatment. At this time point cells accumulate in S/G2/M stages. At 72h of treatment, time point selected for the transcriptomic experiment, the percentage of cells in S/G2/M stages in HeberFERON group exceeded in 10% and 13% with respect to those groups treated with IFNγ and α2b, respectively. Altogether, this ensures an improbable cycling further from Anaphase or ultimately, the impairment of cycle completion and cytokinesis.

Taking into account these elements we propose a general model explaining the distinctive effect of HeberFERON in cell cycle in U-87MG (Figure 10). It is based on the simultaneous activation of the transcriptional factor ATF3 and PKR/EIF2AK2. PKR is significantly up-regulated in IFNα2b and HeberFERON samples, while ATF3 is up-regulated in IFNγ and HeberFERON. That is, the activation of both proteins only occurs in samples treated with HeberFERON. PKR plays an important role in tumor suppression function of p53 (Yoon et al. 2009). It phosphorylates p53 at Ser392 (Cuddihy et al. 1999) and this phosphorylation is important for p53 activation and localization (Castrogiovanni et al. 2018). The second element of the model relies on the role of ATF3 activation on blocking MDM2 degradation of p53 (Yan et al. 2005) and preventing the translocation of p53 to cytoplasm contributing to its tumor-suppressor activity (Lohrum et al. 2001). In Glioblastomas, p53 is known to be located mostly in cytoplasm (Nagpal et al. 2006); its localization in the nucleus is associated to a longer survival of GBM patients (Burton et al. 2002). Sequestration of p53 in the cytoplasm prevents its translocation to the nucleus and presumably avoids its suppressive function; this has also been found in poorly differentiated pediatric neuroblastomas (Moll et al. 1995). Also in primary GBM, the location of p53 wild type in the cytoplasm has been correlated with the expression of Vimentin (Sembritzki et al. 2002). These elements, together with the known functions of ATF3 and PKR, suggest the future study of the location of p53 in cells treated with HeberFERON. Among genes repressed by p53 we have CCNB1 and BIRC5, both genes strongly down-regulated by HeberFERON. Additional mechanism includes p53-mediated increased expression of GADD45 that binds to CDK1 and prevents cyclinB-CDK1 complex formation and G2 arrest (Bai and Zhu 2006). Increase GADD45 and decrease CCNB1 expressions, mediated by p53, can be contributing to the cycle arrest.

**Figure 10:**
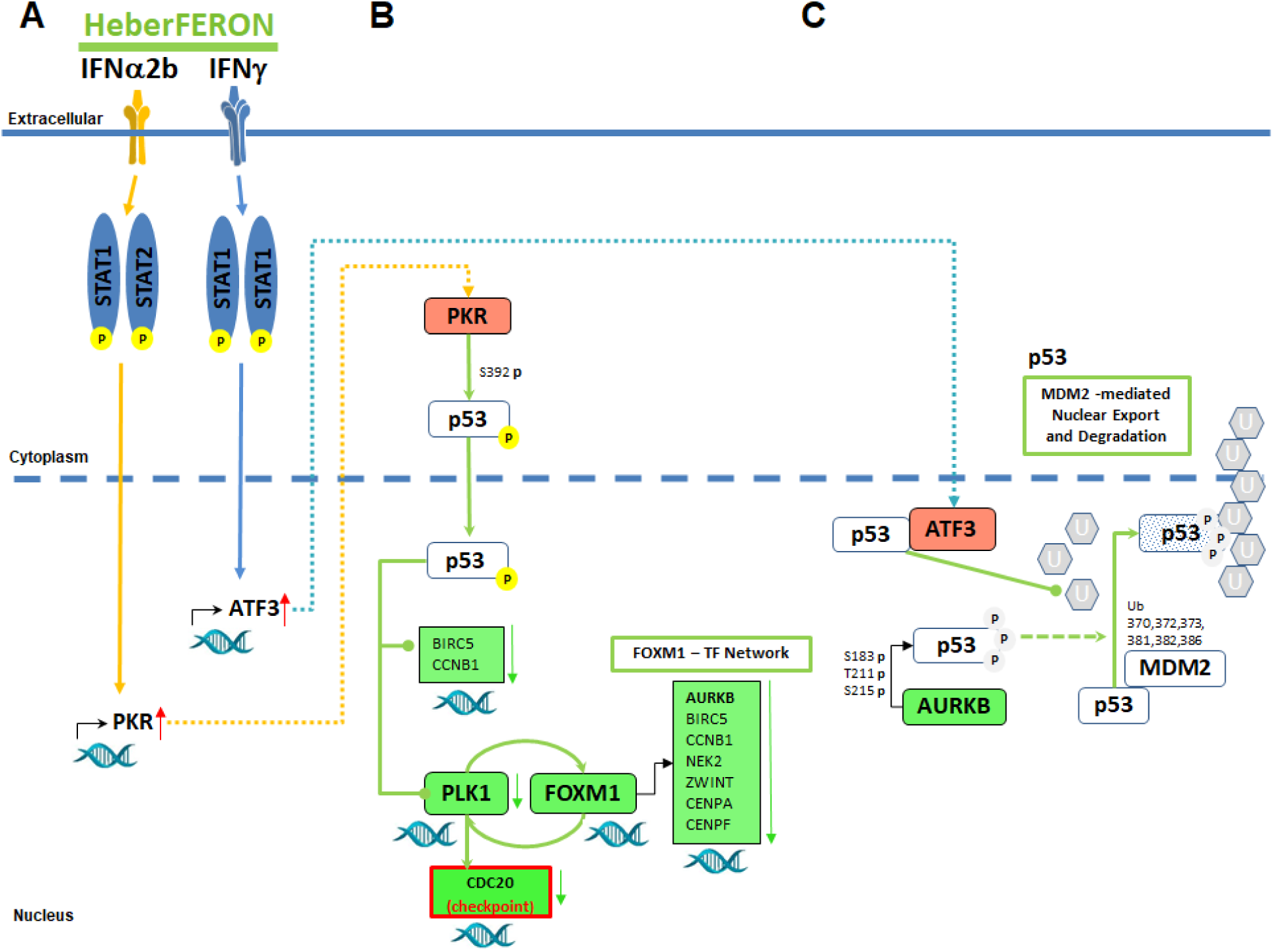
Model proposal of molecular mechanisms involved in the distinctive action of HeberFERON. In this schematic representation, HeberFERON activates PKR and ATF3 simultaneously through the STAT complex. (**A**) IFNα2b and IFNγ induce the up-regulation of PKR and ATF3, respectively. (**B**) PKR phosphorylates p53 at Serine 392, p53 is translocated to the nucleus where it represses the expression of PLK1, consequently FOXM1 phosphorylation is reduced and the FOXM1 transcription network is down-regulated. CDC20, also activated by PLK1, is down-regulated delaying exit from mitosis. (**C**) AURKB, as part of FOXM1 transcription network, is down-regulated and its phosphorylation activity over p53 at residues Ser183, Thr211, and Ser215 is reduced, slowing down p53 degradation through MDM2-mediated ubiquitination. In parallel ATF3 up-regulation increase its association with p53 reducing also MDM2 ubiquitination of p53 and its export from nucleus to cytoplasm. Rounded rectangles filled in red and green means up-regulated and down-regulated gene/proteins. Letter “p” in a yellow circle represent phosphorylated residues, letter “p” in gray circle represent residue with diminished phosphorylation. Letter “U” in a gray hexagon represent ubiquitin protein slowly added to target protein p53.

In the model of Figure 10 we also show FOXM1 network, which is down-regulated by HeberFERON. Phosphorylation of FOXM1 by PLK1 provides a positive feedback loop essential for mitotic progression (Fu et al. 2008). APC/C-CDC20 complex components also need to be phosphorylated by PLK1 to contribute to mitosis stages progression by sequential protein degradation (Sivakumar and Gorbsky 2015). APC/C-CDC20 function will also depend on the mitotic spindle ckeckpoint complex. Additionally, Aurora kinase B is a negative regulator of p53 by phosphorylation on Ser183, Thr211, and Ser215 which contributes to p53 degradation through MDM2-mediated ubiquitination (Gully et al. 2012). All these facts are integrated into the proposed model.

Diagnostic, prognostic or predictive molecular biomarkers are limited in GBM although they could have an impact in OS and personalization of treatments (Kan et al. 2020). The best known is the methylation status of O(6)-methylguanine-DNA-methyltransferase (MGMT) as a predictor of temozolomide (Hegi et al. 2005) and radiation resistance (Rivera et al. 2010). Other has described IFNβ or IFNγ associated gene signatures to predict OS, efficacy of immunotherapy and radiotherapy among glioblastoma patients (Cheng et al. 2021, Ji et al. 2021). Eventually, these genes signatures should be validated in the clinical practice.

Here, we identified a signature of poor prognosis in Proneural subtype of GBM (Duarte et al. 2012). The expression levels of genes belonging to this poor prognosis signature increased with HeberFERON treatment as with IFNα2b treatment. The reformulation of HeberFERON with different IFNα2b and γ proportions could be an alternative to treat this subtype of GBM tumors. This proposal should also be validated.

## Conclusions

As part of this investigation we described the transcriptomic profile of the HeberFERON in comparison to individual IFN treatments. It was important to show that, similar to other IFNs combinations, HeberFERON highly stimulates the transcription of genes involved in antiviral and immune responses. More interesting was to find HeberFERON distinctively targets transcripts encoding proteins that participate in cell cycle events. As part of these events, we found key players in mitotic prometaphase to anaphase stages, spindle checkpoint and proteolytic degradation in mitosis by APC/C-CDC20 which could explain the G2/M arrest and antiproliferation effect over U-87MG. A signaling from the STATs through PKR and ATF3 factors converge to P53 and from this cascade hub the signal propagate to cell cycle and proliferations players with relevance in GBM in a multi-targeted way. These findings support our proposed general mechanistic model and also underscore the use of HeberFERON alone or in combination with chemotherapeutics in the treatment of GBM.

## Supplementary Information

**Additional file 1: Table S1:** Oligonucleotides used for qPCR amplifications.

**Additional file 2: Fig. S1.** Dose-Effect HeberFERON Curves and cell counting.

**Additional file 3: Table S2.** Output of Gene Ontology (GO) analysis for 23 common genes.

**Additional file 4: Fig. S2.** Expression profiles of FOXM1 regulated genes and CDC20 in brain tumors by GlioVis.

**Additional file 5: Fig. S3.** Results of Gene Set Enrichment Analysis (GSEA).

**Additional file 6: Fig. S4.** Cell Cycle gene expression plot.

**Additional file 7: Fig. S5.** Fold changes of genes from a molecular signature associated to poor prognosis in Proneural GBM subtype.

**Authors’ contributions** I.B. conceived the project; D.V-B., J.M. and I.B. designed the microarray experiment; D.V-B. and I.B. obtained and processed the samples for microarray experiment; D.V-B. performed the experimental validation; J.M., R.B. and J.F-C. performed the functional and network analysis; J.M. quality control of expression data and proposed the bioinformatics model; D.V-B., D.P. and L.I.N. followed the microarray experiment service; J.M., R.B. and D.V-B. wrote the manuscript; I.B. and J.F-C. revised the manuscript and suggested changes. All authors read and approved the final manuscript.

## Declarations

### Ethics approval and consent to participate

Not applicable.

## Consent for publication

Not applicable.

## Figures and Tables

### Supplementary Information

**Table S1.**
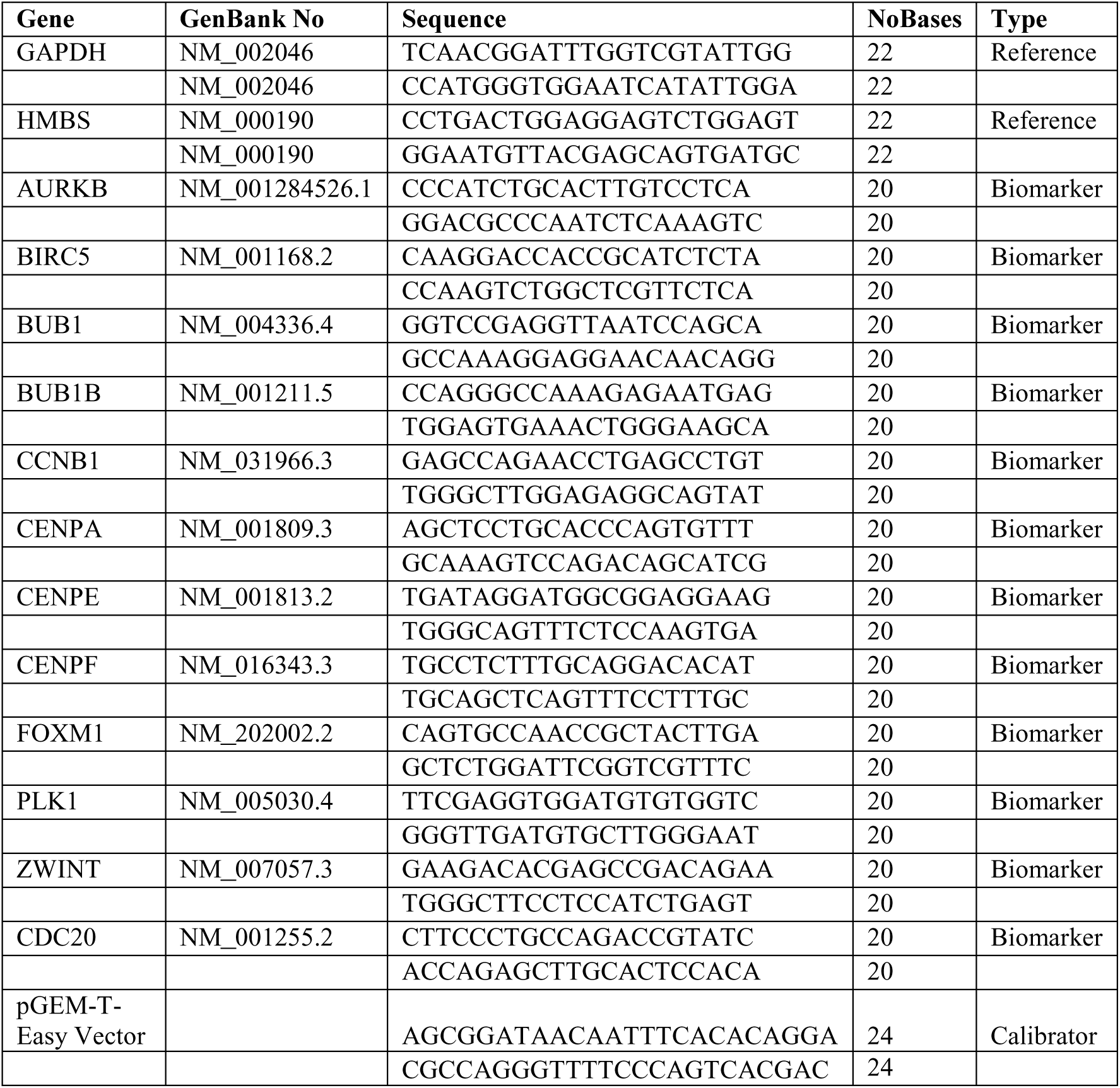
Oligonucleotides used for qPCR amplifications. Information of Gene name, GenBank number (GenBank No), Sequence, number of bases and type (Reference, Biomarker or Calibrator) is provided. All were synthesized in Oligonucleotide Synthesis Group (CIGB, Havana). 10^3^ copies of pGEM-T-Easy Vector (Promega, USA) were used as a calibrator in each run.

**Figure S1:**
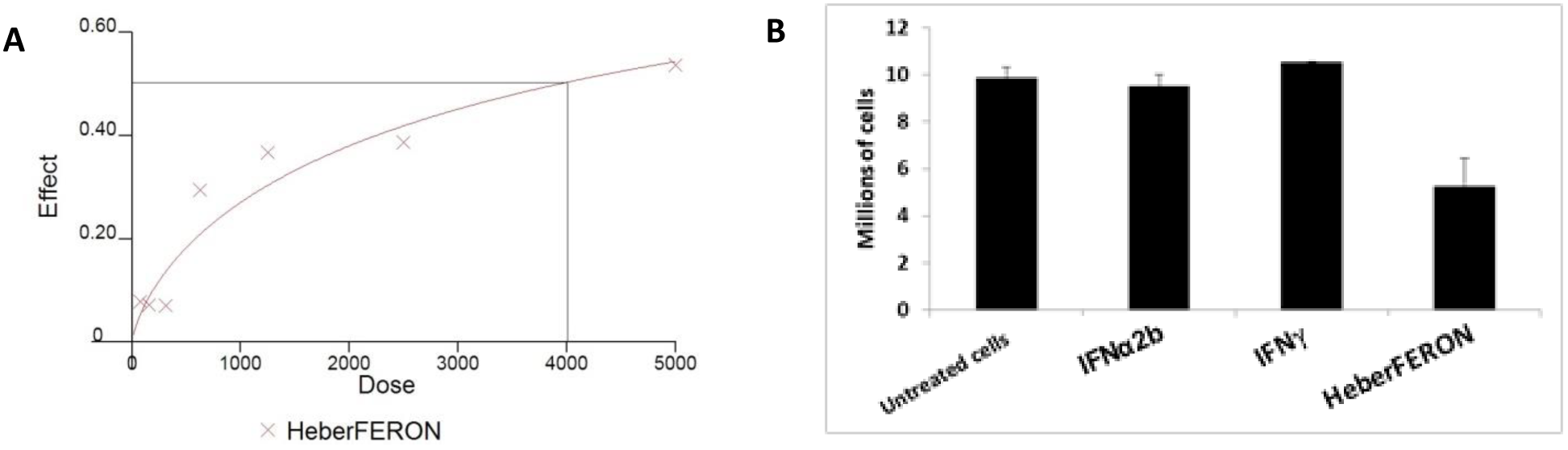
Dose-Effect HeberFERON Curves and cell counting. (A) The cells were treated with HeberFERON at different concentrations for 72 h and cell viability assessed by MTT assay. Relations of concentration in IU/mL (Dose) and Effect (0-0.6, meaning 0-60% of cell proliferation inhibition) is plotted. IC50 is calculated as the concentration to achieve a 50% (0.5) of effect. (B) Cell counting with Trypan blue 0.4% was performed in duplicated cultures treated with HeberFERON at IC50 or their equivalent dose for IFNα2b or IFNγ. Millions of cells average and standard deviation are shown.

**Table S2:** Output of Gene Ontology (GO) analysis for 23 common genes. *Adj p-value<0.05.

| GO Term | Count | % | P-Value | Benjamini |
| --- | --- | --- | --- | --- |
| defense response to virus | 7 | 36.8 | 2.00E-08 | 4.30E-06 |
| innate immune response | 8 | 42.1 | 2.40E-07 | 2.50E-05 |
| antigen processing and presentation of endogenous peptide antigen via MHC class I via ER pathway, TAP-independent | 3 | 15.8 | 1.80E-05 | 1.30E-03 |
| response to interferon-beta | 3 | 15.8 | 3.50E-05 | 1.90E-03 |
| positive regulation of T cell mediated cytotoxicity | 3 | 15.8 | 2.20E-04 | 9.50E-03 |
| type I interferon signaling pathway | 3 | 15.8 | 6.30E-04 | 2.00E-02 |
| negative regulation of viral genome replication | 3 | 15.8 | 6.60E-04 | 2.00E-02 |

**Figure S2:**
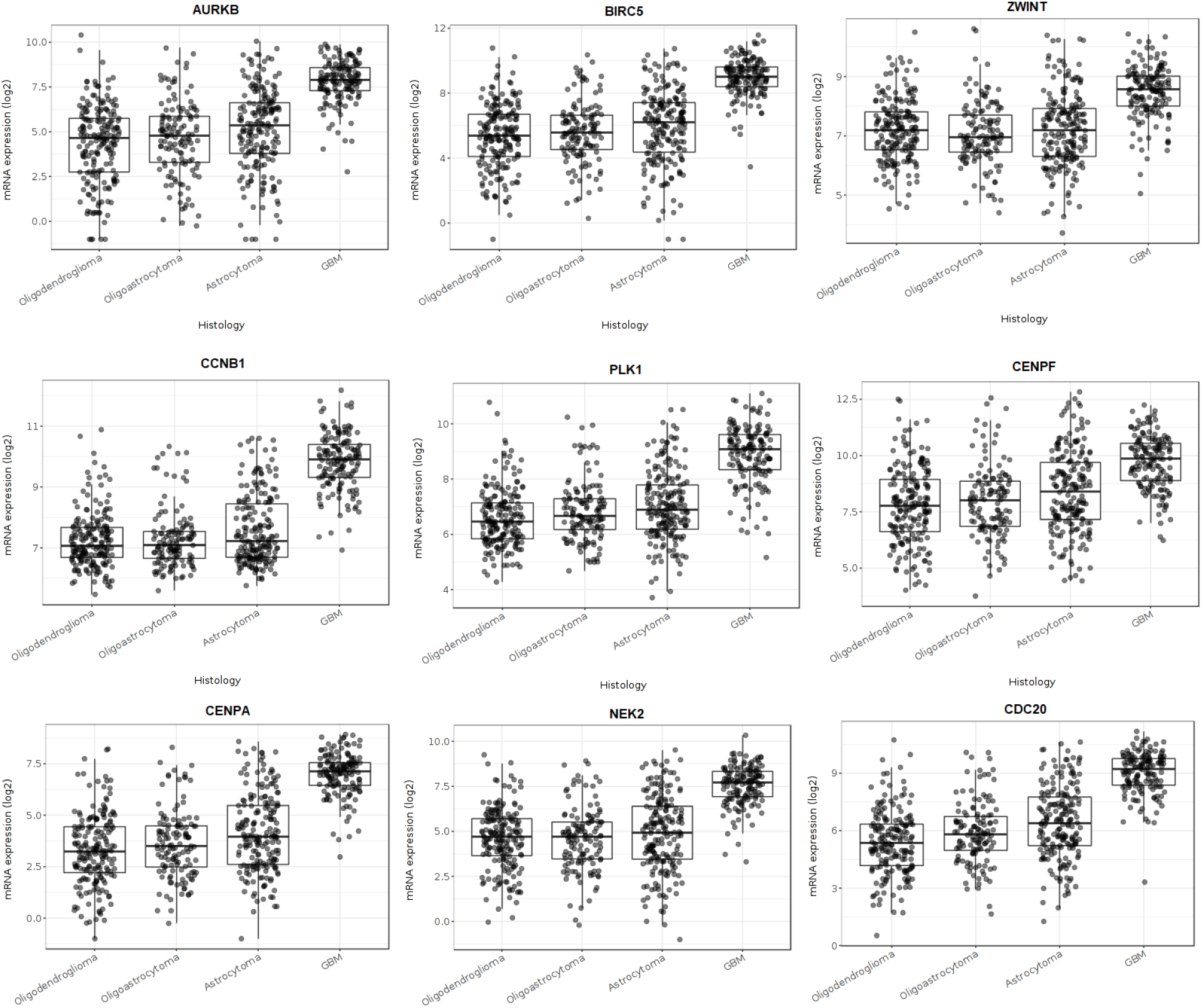
Expression profiles of FOXM1 regulated genes and CDC20 in brain tumors by GlioVis.

**Figure S3:**
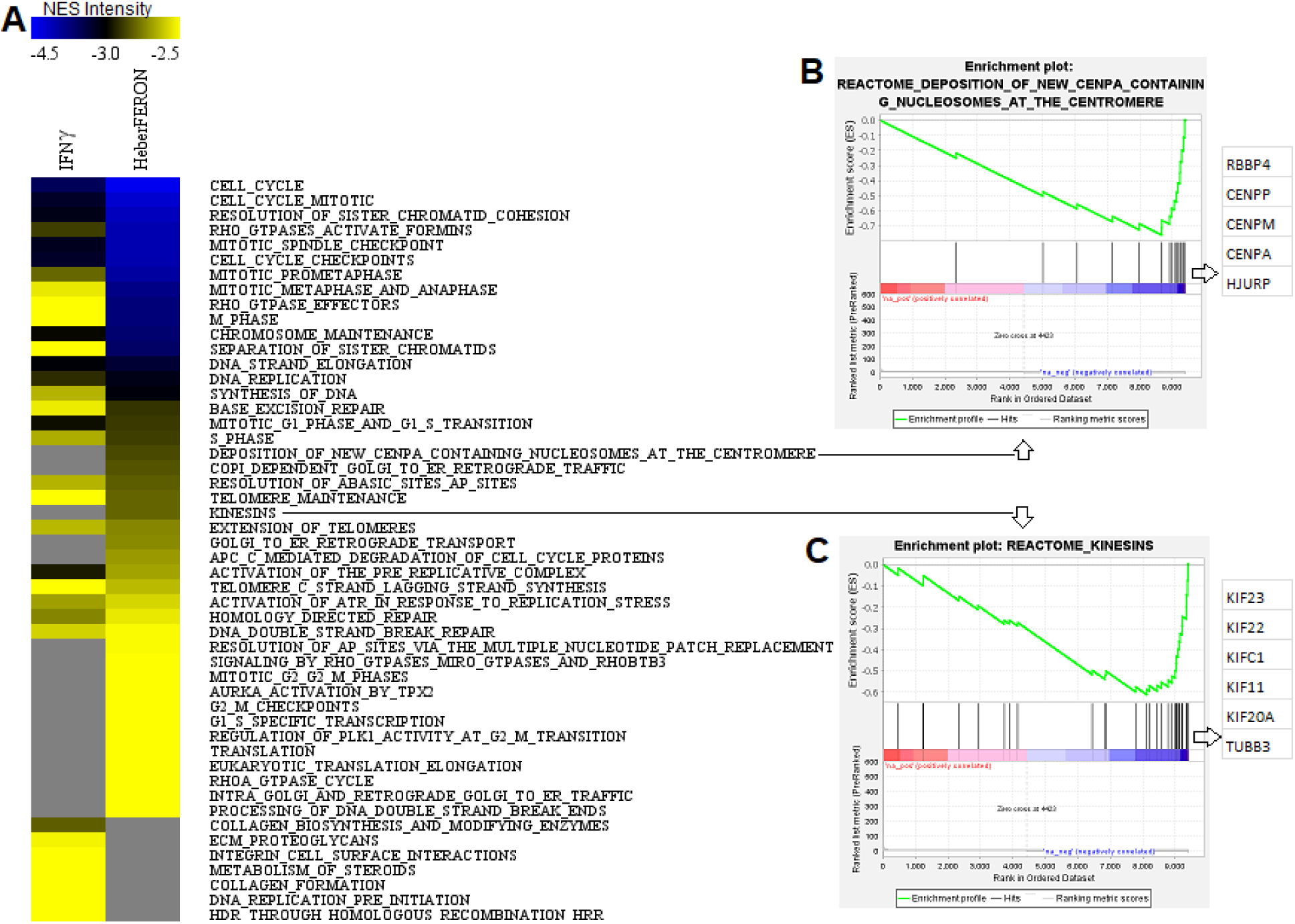
Results of Gene Set Enrichment Analysis (GSEA). (**A**) Heatmap of the Normalized Enrichment Scores (NES) of Reactome pathways resulted from GSEA analysis. The two columns correspond to IFNγ and HeberFERON. Reactome Pathways ordered by NES by HeberFERON. Identification of signaling distinctively regulated (inhibited) by this combination of IFNs α2b and γ. (**B**), (**C**) Two enrichment plots and most relevant genes from Core enrichment set are visualized for Pathways present only under HeberFERON: “Deposition of new CENPA containing nucleosomes at the centromere”(**B**) and Kinesins (**C**). Most relevant genes for the enrichment significance are listed to the right of each Enrichment Plot.

**Figure S4:**
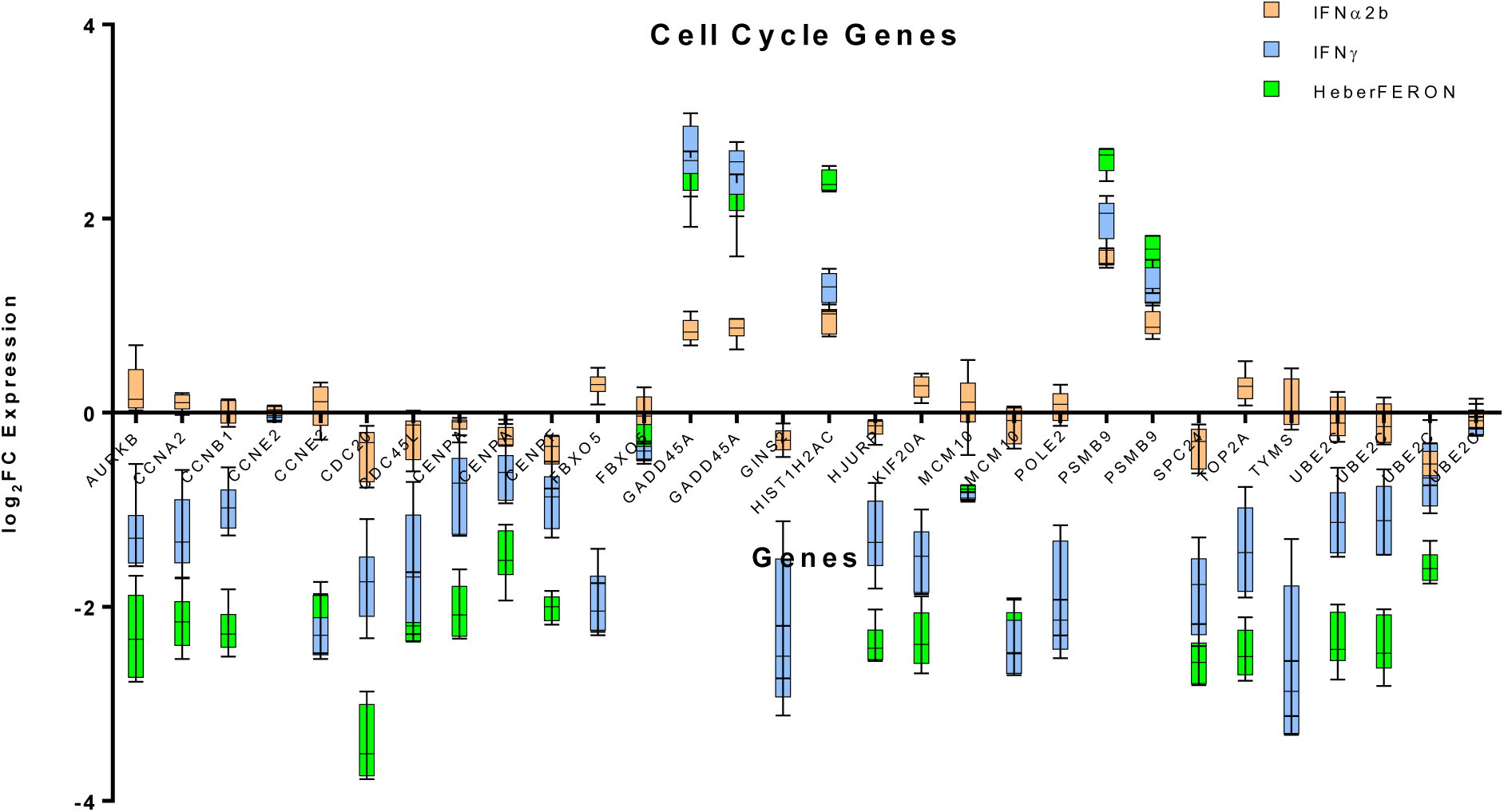
Cell Cycle gene expression plot. log_2_FC expression are plotted. Orange, blue and green color bars correspond to IFNα2b, IFNγ and HeberFERON expression values, respectively.

**Figure S5:**
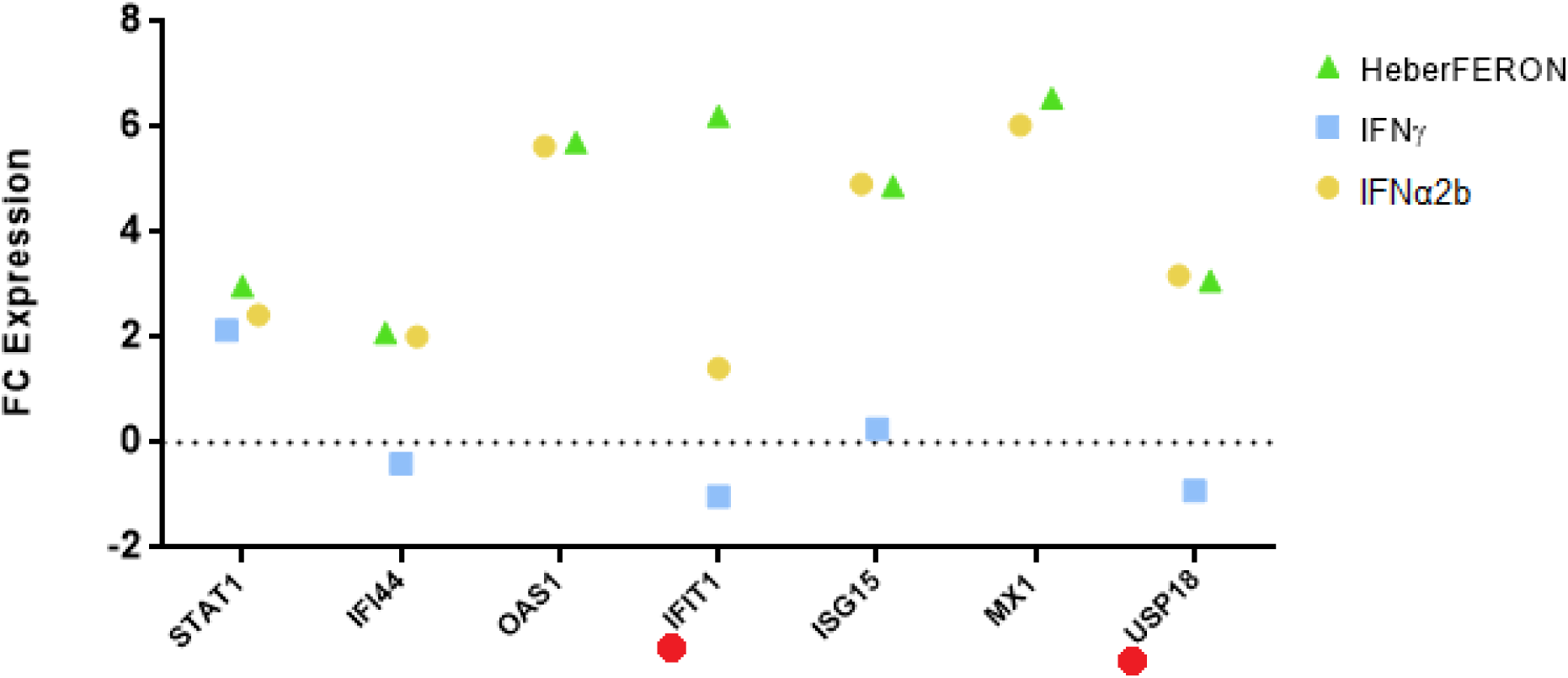
Fold changes of genes from a molecular signature associated to poor prognosis in Proneural GBM subtype. Orange, blue and green represents expression values in samples treated with IFNα2b, IFNγ and HeberFERON, respectively. Red dots highlight genes whose expression is favorable for the disease outcome.

